# Microbiota-stimulated Interleukin-22 regulates brain neurons and protects against stress-induced anxiety

**DOI:** 10.1101/2022.09.16.508224

**Authors:** Anoj Ilanges, Mengyu Xia, Junmei Lu, Lei Chen, Rani Shiao, Changchun Wang, Zheyu Jin, Ru Feng, Qingqing Qi, Han Yi, Jixi Li, Marc Schneeberger, Boxun Lu, Jeffrey M. Friedman, Xiaofei Yu

**Author notes:** These authors contributed equally to this work (Anoj Ilanges, Mengyu Xia, Junmei Lu, Lei Chen). Senior author (Jeffrey M. Friedman). Correspondence (XY).

## Abstract

Psychological stress and its sequelae are a major public health problem. While the immune system has been implicated in the development of stress-related disorders, how the immune signals modulate neural responses to stress is poorly understood. Contrary to our expectations, we found that the immune cytokine Interleukin (IL)-22 is the key mediator of an immune-to-brain pathway that diminishes, rather than amplifies, stress-induced anxiety. We showed that stress induced T_H_17 differentiation and IL-22 production in the intestine following barrier dysfunction and microbiota stimulation. IL-22 then directly signaled to septal neurons in the brain to mitigate anxiety-like behavior. Accordingly, mice treated with exogenous IL-22 showed resilience to chronic stress-induced anxiety disorders. Our study thus reveals a previously-unappreciated immune-to-brain axis that defends against psychological stress, suggesting a potential intervention strategy for stress-related mental diseases.

## Introduction

Animals thrive in constantly changing environments by sensing and mounting coordinated responses to stress. The responses are delicately orchestrated across the nervous system and peripheral organs, which are collectively termed allostasis (McEwen et al., 2015). The dysregulation of this process has been shown to contribute to the development of a broad spectrum of psychological diseases such as anxiety disorders and depression (McEwen *et al*., 2015). However, allostatic responses to stress have been primarily studied from the perspective of the brain (Calhoon and Tye, 2015; McEwen *et al*., 2015), with less known about other organ systems involved.

The immune system, which predominantly monitors and protects the internal integrity of the body, has recently been implicated in allostatic responses to stress. On the one hand, numerous studies suggest that the immune system promotes the development of stress-related psychological diseases, such as anxiety disorders and depression (Glaser and Kiecolt-Glaser, 2005; Haykin and Rolls, 2021; Huh and Veiga-Fernandes, 2020; Menard et al., 2017; Pape et al., 2019; Salvador et al., 2021). For example, inflammation and immune dysregulation have been observed in patients with stress-related diseases. T lymphocytes, an important arm of the immune system, have also been reported to induce anxiety-like behavior by producing metabolic or cytokine signals, particularly when pathologically activated (Alves de Lima et al., 2020; Choi et al., 2016; Fan et al., 2019; Miyajima et al., 2017). On the other hand, fewer studies have also indicated that lymphocytes, including CD4^+^ T cells, could confer stress resilience and ameliorate stress-induced psychological disorders (Brachman et al., 2015; Cohen et al., 2006; Rattazzi et al., 2013; Scheinert et al., 2016), but the underlying mechanisms remain elusive. Therefore, it is likely that the immune system is key to the balanced allostatic response to stress (Medzhitov, 2021), which may also exacerbate psychological disorders if dysregulated. Given the immune sensing of resident microbiota that has been shown to influence host behavior (Li et al., 2022; Morais et al., 2021; Needham et al., 2022; Veiga-Fernandes and Mucida, 2016), the immune system may integrate microbial danger signals to instruct the host response to stress. Therefore, it is critical to understand how the immune system responds to stress and how the immune response integrates with neural pathways during stress, which are yet to be elucidated.

## Results

### IL-22 is sufficient and necessary to suppress anxiety in response to stress

To investigate whether and how the immune system participates in the allostatic response to stress, we studied stress-induced anxiety in mice lacking the T cell receptor β subunit (*Tcrb*^*-/-*^) (Mombaerts et al., 1992). *Tcrb*^*-/-*^ mice have targeted deficiency in *αβ* T cells and develop an imbalanced immune system but are generally healthy when maintained in SPF facilities. We reasoned that *Tcrb*^*-/-*^ mice could represent an immune imbalance state that might alter immune-to-brain signaling while minimizing potential confounding effects of physical illness. We assessed allostatic responses to stress by testing for anxiety-like behavior using the classical open field test (OFT; more time spent in the center of the arena is indicative of reduced anxiety) and elevated plus maze (EPM; more time spent in the open arms of the arena is indicative of reduced anxiety) (Calhoon and Tye, 2015). Importantly, at the resting state, wild-type (WT) and *Tcrb*^*-/-*^ mice spent similar amounts of time exploring the center of the OFT arena (Figure 1A and 1B) and the open arms of the EPM maze (Figure S1A and S1B). As an internal control, wild-type (WT) and *Tcrb*^*-/-*^ mice traveled similar distances (Figure 1A and 1B), indicating comparable locomotor activities. Thus, *Tcrb*^*-/-*^ mice exhibited comparable anxiety-like behavior as WT mice at baseline, making them an ideal discovery model to study immune-brain communications in response to stress.

**Figure 1.**
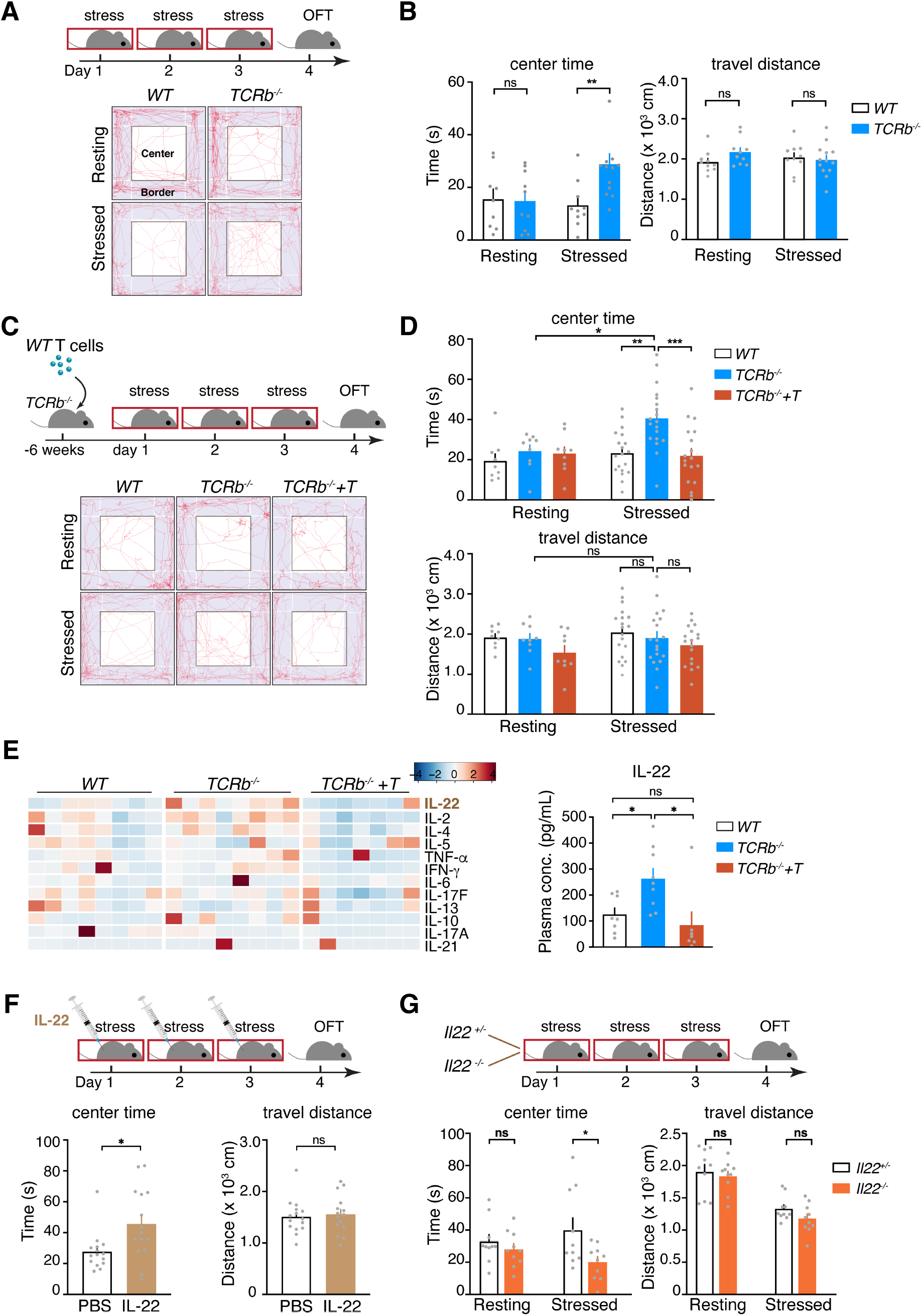
IL-22 reduces mouse anxiety in response to stress. (**A**,**B**) Wild-type (WT) and T cell-deficient *Tcrb*^*-/-*^ mice were subjected to repeated mild restraint stress (30-minutes restraint/day for 3 days) and examined for anxiety-like behavior with the Open Field Test (OFT). (A) Illustration of the experimental procedure and representative track plots. (B) Quantification of mouse anxiety (center time) and locomotor activity (travel distance, as an internal control). n=9, 10, 10, 13 for WT-resting, *Tcrb*^*-/-*^-resting, WT-stressed, and *Tcrb*^*-/-*^-stressed, respectively. (**C**,**D**) WT, *Tcrb*^*-/-*^ and T cell-rescued *Tcrb*^*-/-*^ (*Tcrb*^*-/-*^*+T*) mice were stressed and tested for anxiety. (C) Illustration of the experimental procedure and representative track. (D) Quantification of mouse anxiety and locomotor activity. n=9, 9, 9, 19, 19, 18 for WT-resting, *Tcrb*^*-/-*^-resting, *Tcrb*^*-/-*^*+T*-resting, WT-stressed, *Tcrb*^*-/-*^-stressed and *Tcrb*^*-/-*^*+T*-stressed, respectively. **(E)** Circulating cytokine levels in WT, *Tcrb*^*-/-*^ and *Tcrb*^*-/-*^*+T* mice (heatmap), with IL-22 concentrations shown in a bar plot. n=8, 8, 7 for WT, *Tcrb*^*-/-*^ and *Tcrb*^*-/-*^*+T* mice, respectively. **(F)** WT mice were treated with recombinant mouse IL-22 and tested for anxiety-like behavior in response to stress. n=15 for both PBS and IL-22-treatment groups. **(G)** *Il22*^*-/-*^ and *Il22*^*+/-*^ control mice were analyzed for anxiety-like behavior. n=10 for both groups. Data were pooled from or representative of 2-3 independent experiments and shown as Means ± SEMs. Statistical analysis was performed with 2-way ANOVA with Sidak’s multiple comparisons (B,D,G), 1-way ANOVA with Tukey’s multiple comparisons (E), and Unpaired t-test (F). *P < 0.05; **P < 0.01; ***P < 0.001; ns, not significant.

We then examined stress-induced anxiety by subjecting both groups to repeated mild stress, which consists of a daily 30-minute period of restraint for 3 consecutive days. This paradigm introduces stress but is not sufficient to induce mood disorders such as anxiety disorders and depression that usually require 2 hours of restraint per day for 2-3 weeks (Buynitsky and Mostofsky, 2009). The time frame of our treatment paradigm is also long enough to allow early immune responses to occur. Compared to similarly stressed WT mice, *Tcrb*^*-/-*^ mice spent more time exploring the center of the OFT arena and the open arms of the EPM maze, while traveling similar distances. Therefore, *Tcrb*^*-/-*^ mice showed reduced anxiety compared to WT mice after mild stress exposure (Figure 1A, 1B, S1A and S1B).

We next tested whether this difference could be attributed to T cells by transplanting WT T cells into *Tcrb*^*-/-*^ (*Tcrb*^*-/-*^*+T*) mice (Figure 1C, S1C and S1D). We found that introducing normal T cells increased the anxiety-like behavior of *Tcrb*^*-/-*^ mice after stress, equivalent to that observed in WT mice (Figure 1C, 1D, S1E and S1F). Given that *αβ* T cells are critical for maintaining immune balance, these data collectively suggested that a T cell-regulated immune signal modulates anxiety-like behavior in response to psychological stress.

To identify the putative signal(s), we profiled peripheral cytokines, the primary immune signals mediating communications between the immune system and other organ systems such as the brain (Glaser and Kiecolt-Glaser, 2005; Salvador *et al*., 2021), in stressed WT, *Tcrb*^*-/-*^ and *Tcrb*^*-/-*^*+T* mice (Figure 1E). Coupled with behavior data, we found that there was a significant inverse correlation between IL-22 levels and anxiety-like behavior: IL-22 concentrations were significantly higher in *Tcrb*^*-/-*^ mice than in WT mice, with restoration to WT levels in *Tcrb*^*-/-*^*+T* mice (Figure 1E). No clear correlation was detected for all other cytokines examined, including IFN-*γ* and IL-17A which have been previously shown to directly signal to CNS neurons to modify social and anxiety-like behavior, respectively (Alves de Lima *et al*., 2020; Choi *et al*., 2016; Filiano et al., 2016; Reed et al., 2020).

To test whether IL-22 was sufficient to exert an anxiolytic effect, we intraperitoneally (i.p.) injected WT mice with recombinant IL-22 and subjected them to mild restraint stress (Figure 1F). IL-22 injection significantly reduced stress-induced anxiety-like behavior, recapitulating the phenotype of *Tcrb*^*-/-*^ mice with higher IL-22. We then examined whether IL-22 was necessary for maintaining proper anxiety in response to stress. While IL-22-deficient (*IL-22*^*-/-*^) and *IL-22*^*+/-*^ control mice showed no difference in anxiety at baseline, *IL-22*^*-/-*^ mice expressed increased anxiety after stress (Figure 1G). Together, our data showed that IL-22 is both necessary and sufficient to regulate anxiety in response to stress. We next set out to define its site of action.

### The septal area of the brain is an important interfacing point for IL-22 to suppress anxiety

Stress activates multiple brain areas to elicit anxiety (Calhoon and Tye, 2015), so we hypothesized that IL-22 might modulate these stress-activated neural pathways. Since *Tcrb*^*-/-*^ mice are intrinsically high in IL-22, we mapped brain-wide differential neural activities between WT and *Tcrb*^*-/-*^ mice by examining the expression of the immediate-early gene cFOS with whole-mount brain clearing and immunostaining (iDISCO) (Renier et al., 2014). We performed a time-course analysis to fully capture neural activities in response to stress, which consists of 4 time points: baseline (Resting), two hours after the start of restraint stress (2h), one day after stress (1-day), and two hours after the OFT performed one day after stress (1-day OFT) (Figure 2A). WT and *Tcrb*^*-/-*^ mice showed similar neuronal activation profiles at baseline (Figure S2A and S2B), in line with comparable anxiety-like behavior at the resting state. In contrast, WT and *Tcrb*^*-/-*^ mice showed substantial differences in neuronal activation after restraint stress (2h, 1-day, 1-day OFT) (Figure S2A), in brain regions involved in sensation and emotion processing such as primary somatosensory areas (SSp), cortical amygdalar area (COA), bed nuclei of the stria terminalis (BST) and lateral septal complex (LSX) at one or more post-stress timepoints (Figure S2B).

**Figure 2.**
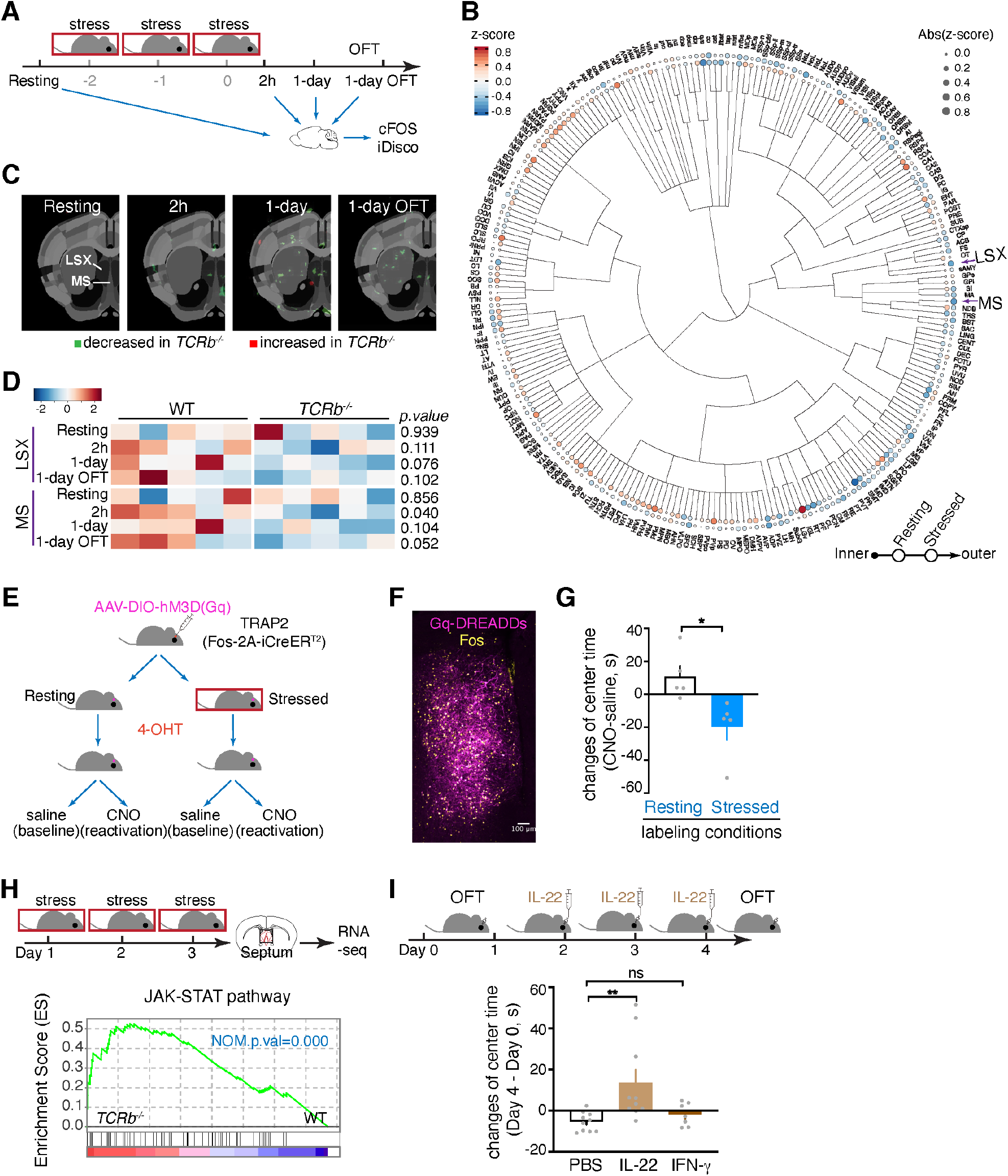
The brain septal area is an important interfacing point for IL-22 to suppress anxiety. (**A-D**) Unbiased time course profiling of neuronal activation in WT and *Tcrb*^*-/-*^ mice was performed by examining the expression of cFOS in the whole brain with iDISCO whole-mount imaging. (A) Illustration of the experimental design and timepoints examined: baseline (Resting), two hours after the final restraint stress (2h), one day after the final restraint stress (1-day), and after exposure to OFT one day after the final restraint stress (1-day OFT). (B) Differential activation of brain regions by stress in WT and *Tcrb*^*-/-*^ mice. The inner dots show baseline (Resting) z scores of each brain region and the outer dots show aggregate z scores of post-stress timepoints (2h, 1-day, 1-day OFT). Blue and red indicate negative and positive z scores, respectively, corresponding to decreased and increased neuronal activation in *Tcrb*^*-/-*^ mice compared to WT mice. The size of dots and intensity of colors reflect the absolute values of z scores. The tree structure shows the hierarchical organization of brain regions. (C) Statistical analysis of neuronal activation in the septal area at the mapped pixel level. Green and red indicate significant decrease (*p*<0.01) and increase (p<0.01) in *Tcrb*^*-/-*^ mice. (D) Heatmap analysis of neuronal activation in the septal area at the region level. n=5 for all groups. (**E-G**) Reactivation of stress-activated septal neurons to examine their sufficiency to drive anxiety-like behavior. (E) Design of the reactivation experiment. Stress-activated septal neurons were labeled with the activating hM3D(Gq) DREADD receptor in TRAP2 mice and reactivated by CNO. (F) Representative images showing labeling of activated (cFOS^+^, yellow) septal neurons by hM3D(Gq) (magenta) in stressed mice. (G) Changes of center time of individual mice injected with CNO and saline (baseline) were calculated to reflect the anxiety-driving effects of septal neurons. n=5 for both groups. **(H)** RNA-seq analysis of septal tissues of WT and *Tcrb*^*-/-*^ mice. GSEA analysis was performed to compare the enrichment of JAK-STAT pathway genes in the septum. n=2 for both groups. **(I)** WT mice were stereotactically injected with IL-22 into the septum and analyzed for anxiety-like behavior. Changes in the center time of individual mice before and after IL-22 injection were calculated and depicted. IFN-*γ* was injected as a control. n=10, 10, and 7 for PBS, IL-22, and IFN-*γ* groups, respectively. Data were pooled from 2 independent experiments for (I) and presented as Means ± SEMs. Statistical analysis was performed with ClearMap (B,C,D), Unpaired t-test (G), GSEA (H), and 1-way ANOVA with Dunnett’s multiple comparisons (**I**). *P < 0.05; **P < 0.01; ns, not significant.

To pinpoint the specific brain region(s) responsible for attenuated anxiety in *Tcrb*^*-/-*^ mice after stress, we focused on those regions that showed no difference at baseline (Resting) between WT and *Tcrb*^*-/-*^ mice but showed consistent differences at all three post-stress timepoints (2h, 1-day, and 1-day OFT). By summarizing the post-stress *Tcrb*^*-/-*^-to-WT differences into aggregate z scores, we found that the medial septal nucleus (MS), anterodorsal preoptic nucleus (ADP), ventral medial nucleus of the thalamus (VM), and lateral septal nucleus (LSX) showed most decreased neural activation while the tuberomammillary nucleus (TM) and spinal nucleus of the trigeminal (caudal part) (SPVC) showed most increased neural activation in stressed *Tcrb*^*-/-*^ mice compared to WT mice (Figure 2B). MS and LSX were of particular interest as they are subregions of the septal area that is involved in the anxiety regulation (Calhoon and Tye, 2015). Significant reduction of neural activation in *Tcrb*^*-/-*^ mice was detected in MS and LSX at both mapped pixel and region levels after stress with no difference at baseline (Figure 2B, 2C and 2D). The whole septal area expressed 17.1%, 64.6%, and 32.6% fewer activated neurons in *Tcrb*^*-/-*^ mice than in WT mice at 2h, 1-day, and 1-day OFT, respectively (Figure S2C). Together, these data demonstrated reduced neural activation in the septal area of *Tcrb*^*-/-*^ mice in response to stress (2h) that persisted to the next day (1-day) and after exposure to OFT (1-day OFT).

To test whether stress-activated septal neurons were sufficient to drive anxiety-like behavior, we selectively labeled stress-activated septal neurons with the neuron-activating hM3D(Gq) DREADD receptor (Figure 2E and 2F). This was achieved by stereotactic injection of an AAV encoding a Cre-dependent hM3D(Gq) into the septal area of TRAP2 (*Fos*^*2A-iCreERT2*^) mice (DeNardo et al., 2019), followed by administration of 4-hydroxytamoxifen (4-OHT) under resting and stressed conditions, which enabled hM3D(Gq) expression and labeling of active neurons under either condition. Reactivation of septal neurons labeled at the resting condition with clozapine N-oxide (CNO) did not affect anxiety in mice (Figure 2G). In contrast, reactivation of septal neurons labeled under the stressed condition significantly increased anxiety-like behavior. Thus, stress-activated septal neurons were sufficient to drive anxiety-like behavior, indicating that reduced septal neuron activation by stress in *Tcrb*^*-/-*^ mice may underlie their decreased anxiety-like behavior.

To delineate the mechanisms of reduced septal neuron activation by stress in *Tcrb*^*-/-*^ mice, we performed RNA-seq analysis of septal tissues from stressed WT and *Tcrb*^*-/-*^ mice. Analysis of the expression of cell type-specific markers failed to reveal any significant differences in the overall cellular composition of the septal area (Figure S2D). Nevertheless, Gene Set Enrichment Analysis (GSEA) revealed that a 206-gene stress-induced signature derived from our previous work (Azevedo et al., 2020) was significantly reduced in *Tcrb*^*-/-*^ mice compared to *WT* mice (Figure S2E), in line with reduced neuronal activity in stressed *Tcrb*^*-/-*^ mice. Consistent with the immune imbalance in *Tcrb*^*-/-*^ mice, several immune signaling pathways were highly enriched in *Tcrb*^*-/-*^ mice, with the most significant alterations in the JAK-STAT pathway which mediates the signaling of numerous immune cytokines including IL-22 (Figure 2H and S2F). Indeed, a recent report has confirmed that IL-22 activates the JAK-STAT pathway in an immortalized mouse hippocampal cell line (Lee et al., 2022). Because the septal area is anatomically adjacent to the recently characterized plexus vascular barrier (GVB) that is permissive to large molecules (Carloni et al., 2021), our data raised the possibility that IL-22 might directly signal to neurons in the septal area to modulate anxiety-like behavior.

To test this hypothesis, we implanted cannulas locally into the septal area and assayed anxiety-like behavior after injecting saline or IL-22 (Figure 2I). Similar to systemic administration (Figure 1F), local injection of IL-22 into the septum significantly decreased anxiety-like behavior in mice, suggesting that septal IL-22 signaling is sufficient to suppress anxiety. In contrast, local injection of IFN-*γ*, another cytokine signaling through JAK-STAT failed to induce an effect on anxiety (Figure 2I). Together, our data suggest that the septal area regulates anxiety in response to stress and could be a direct target of IL-22.

### IL-22 suppresses septal neuron activation

We next sought to identify the cellular target of IL-22 in the septal area. IL-22 signals through a dimeric receptor consisting of a selective IL22RA1 subunit and a shared IL10RB subunit (Ouyang and O’Garra, 2019). We examined *Il22ra1* expression in the septum by single-molecule RNA *in situ* hybridization (RNAscope) and found that it was largely co-expressed with the pan-neuronal marker *Rbfox3* (encoding NeuN) (Figure 3A). To further characterize IL-22 receptor-expressing neurons in the septum, we purified NeuN^+^ nuclei by fluorescence-activated cell sorting (FACS) and performed single-nuclei RNA-sequencing (snRNA-Seq) (Figure S3A and S3B). Neurons in the septum show distinct gene expression profiles and cluster separately from neurons of the adjacent cortical/hippocampal and striatal areas (Figure S3C). A total of 6735 septal neurons were further partitioned into 17 clusters, with each cluster defining a distinct cell type. Fourteen clusters express *vGat*(*Slc32a1*)/*Gad1*/*Gad2* and are inhibitory and three clusters express *vGlut2*(*Slc17a6*)/*vGlut1*(*Slc17a7)* and are excitatory (Figure 3B, 3C, S3D and S3E). Though snRNA-seq is limited by the low amount of mRNA in the nucleus (Slyper et al., 2020) and may thus underestimate gene expression, we successfully detected the expression of both IL-22 receptor subunits, *Il22ra1* and *Il10rb*, in septal neurons (Figure 3D and S3E). Simultaneous expression of both receptor subunits was also confirmed at the individual nucleus level (Figure 3E). 79.5% of IL-22 receptor-expressing neurons are GABAergic, including clusters in1, in3, in5, in6, in8, in10, in14, and 20.5% are glutamatergic including cluster ex2 (Figure 3E). Therefore, our data demonstrated that the IL-22 receptor apparatus is expressed by septal neurons.

**Figure 3.**
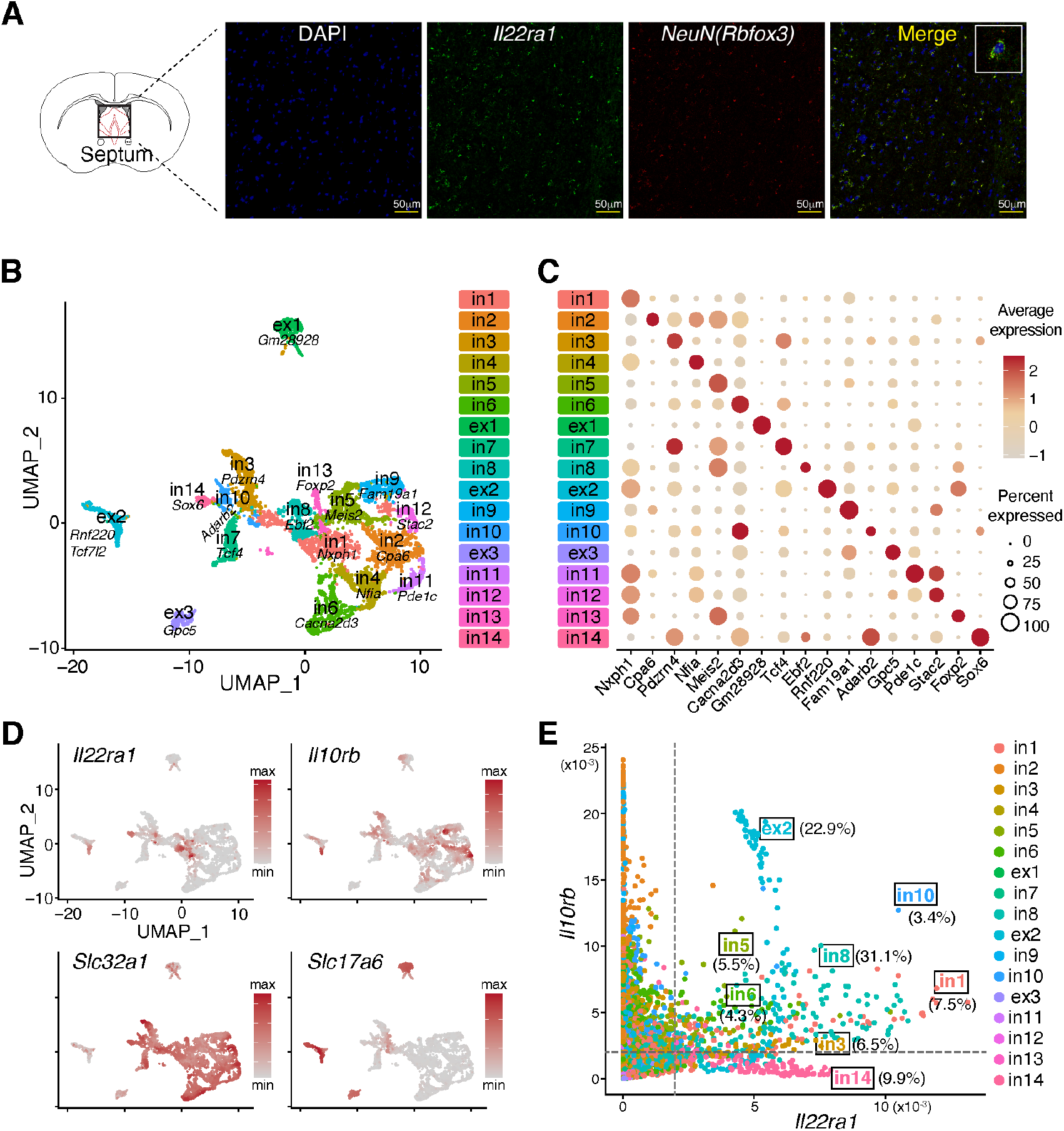
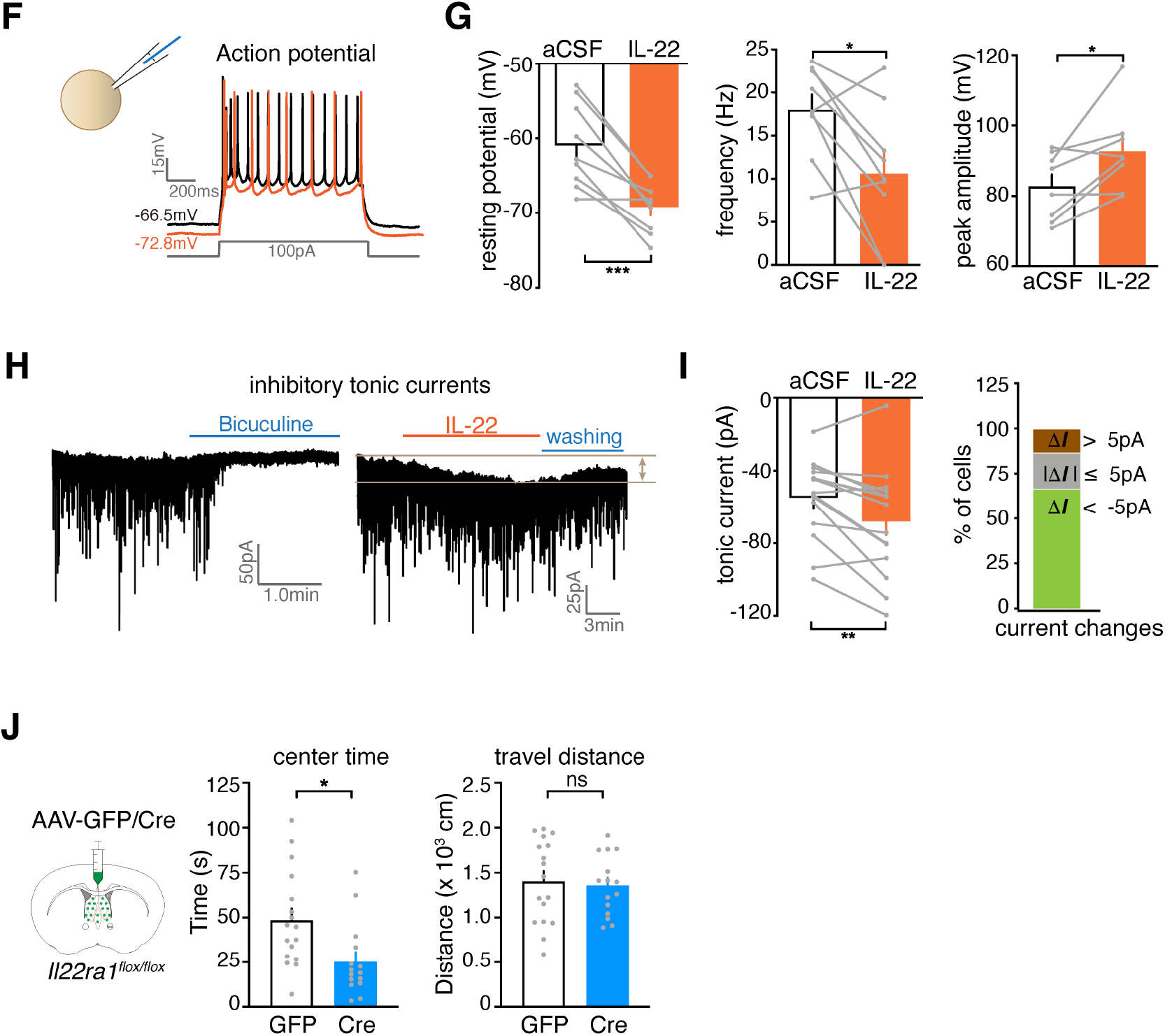
IL-22 inhibits septal neuron activation and reduces anxiety. **(A)** Detection of the IL-22 receptor *Il22ra1* expression by RNAscope. Representative staining of DNA (DAPI, blue), *Il22ra1* (green), the pan-neuronal marker *NeuN* (*Rbfox3*, red), and the merged image are shown. (**B-E**) snRNA-seq analysis of septal neurons. (B) UMAP embedding of single-cell profiles (dots) for the septal area, colored by cluster membership. (C) Dotplot showing top marker genes for each septal neuron cluster. (D) Expression of *Il22ra1, Il10rb*, vGat (*Slc32a1*), and vGlut2 (*Slc17a6*) in septal neurons. (E) Co-expression of *Il22ra1* and *Il10rb* in individual neurons. Ratios of neurons expressing both receptor subunits in each cluster are shown. (**F-I**) Electrophysiological analysis of septal neurons responding to IL-22. (F) Representative action potential recording from septal neurons. (G) Summary of the membrane potential and the frequency and peak amplitude of the action potential of septal neurons treated with buffer (aCSF) or IL-22. (H) Example current trace of mIPSCs when septal neurons were treated with bicuculline (a blocker, left) and IL-22 (right). (I) Quantification of tonic current shift and the proportion of tonic current shift when septal neurons were treated with IL-22. (**J**) *Il22ra1*^*flox/flox*^ mice were stereotactically injected with a *Syn-Cre-GFP* AAV to locally delete *Il22ra1* in septal neurons and analyzed for anxiety-like behavior. n=17, 15 for GFP (control) and Cre groups, respectively. Data were pooled from 2 independent experiments for (J). Data are presented as Means ± SEMs. Statistical analysis was performed with Paired t-test (G,I) and Unpaired t-test (J). *P < 0.05; **P < 0.01; ns, not significant.

The expression of IL-22 receptor subunits in septal neurons suggested the possibility for neurons to respond directly to IL-22. To test this hypothesis, we performed whole-cell patch-clamp recordings on acutely prepared septal slices in the presence or absence of IL-22 (Figure 3F). IL-22 significantly decreased the resting potential of septal neurons, and concomitantly reduced firing frequencies and increased peak amplitudes of action potentials (Figure 3F and 3G). IL-22 treatment also led to an increase of inhibitory tonic currents, with larger than 5 pA inward currents detected in 66.7% of examined neurons (Figure 3H and 3I). These data showed direct suppressive effects of IL-22 on septal neurons. Interestingly, a recent report showed that an immortalized mouse hippocampal cell line could also respond to IL-22 (Lee *et al*., 2022).

We next sought to determine if alterations of IL-22 signaling in septal neurons could affect anxiety-like behavior. Because our snRNA-seq data indicated most IL-22 receptor-expressing neurons were GABAergic inhibitory neurons, we crossed *Il22ra1*^*flox/flox*^ mice with *vGat-Cre* mice to delete IL22RA1 from GABAergic neurons (*Il22ra1*^*ΔvGat*^) (Figure S3F). We found that *Il22ra1*^*ΔvGat*^ mice displayed increased anxiety compared to control mice, indicating that IL-22 signaling in GABAergic neurons is critical for maintaining proper anxiety at baseline. We next ablated the IL-22 receptor specifically in septal neurons by stereotactic injection of a *Syn-Cre-GFP* AAV into the septum of *Il22ra1*^*flox/flox*^ mice (Figure 3J). Compared to control-injected mice (*Syn-GFP* AAV), *Cre*-injected *Il22ra1*-deleted mice showed increased anxiety, suggesting IL-22 signaling in septal neurons mediated an anxiolytic effect. In aggregate, these data showed that IL-22 reduces anxiety by suppressing the activity of septal neurons that express its receptor.

### IL-22 is induced by stress via an IL-1β-T_H_17 pathway

To understand the physiological role of IL-22 in response to stress, we investigated the dynamics of IL-22 during stress. IL-22 is known to maintain host barrier integrity against microbial insults and the intestine is a major source of IL-22, where abundant microbes reside (Ahlfors et al., 2014; Keir et al., 2020; Shaler et al., 2021). Since psychological stress increases intestinal permeability (termed “leaky gut”), exposing host tissues to luminal contents, such as microbiota (Shaler *et al*., 2021; Soderholm and Perdue, 2001), we reasoned that IL-22 could be induced by stress in the gut. Supporting our hypothesis, we observed increased gut permeability to orally gavaged Dextran-FITC and microbial exposure in mice treated with our repeated mild stress paradigm (Figure 4A). Accordantly, IL-22 increased significantly in the blood after stress (Figure 4B), consistent with a recent report (Shaler *et al*., 2021). We then quantified major IL-22-producing cells (T_H_17, ILC3, and *γδ* T cells) in the gut (Ahlfors *et al*., 2014). While no difference was observed in ILC3 cells and *γδ* T cells before and after stress, IL-22-producing T_H_17 cells showed significant increases in the frequency and absolute number after stress (Figure 4C, S4A and S4B). Thus, stress induces T_H_17 cells that produce IL-22 in the gut.

**Figure 4.**
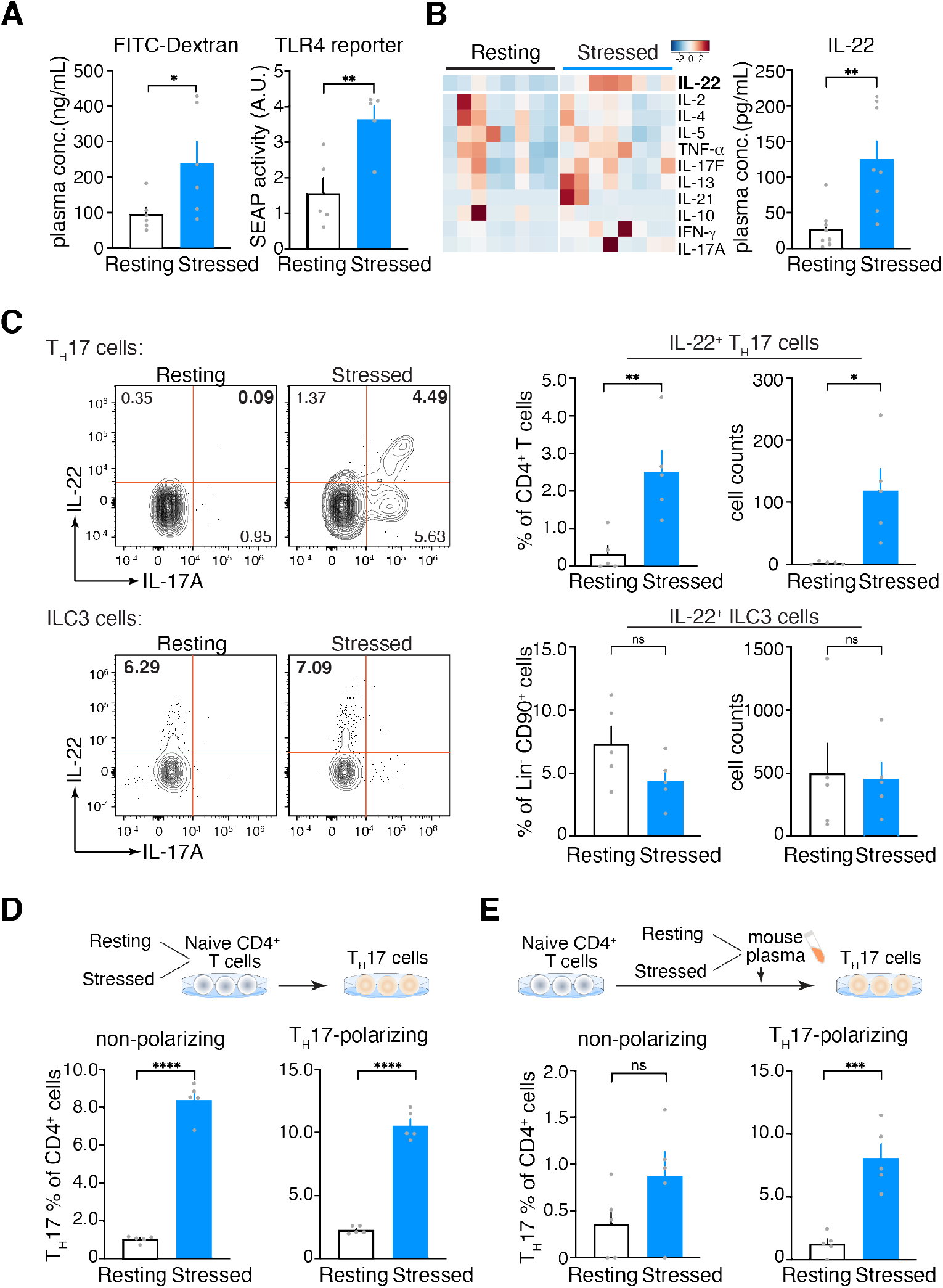
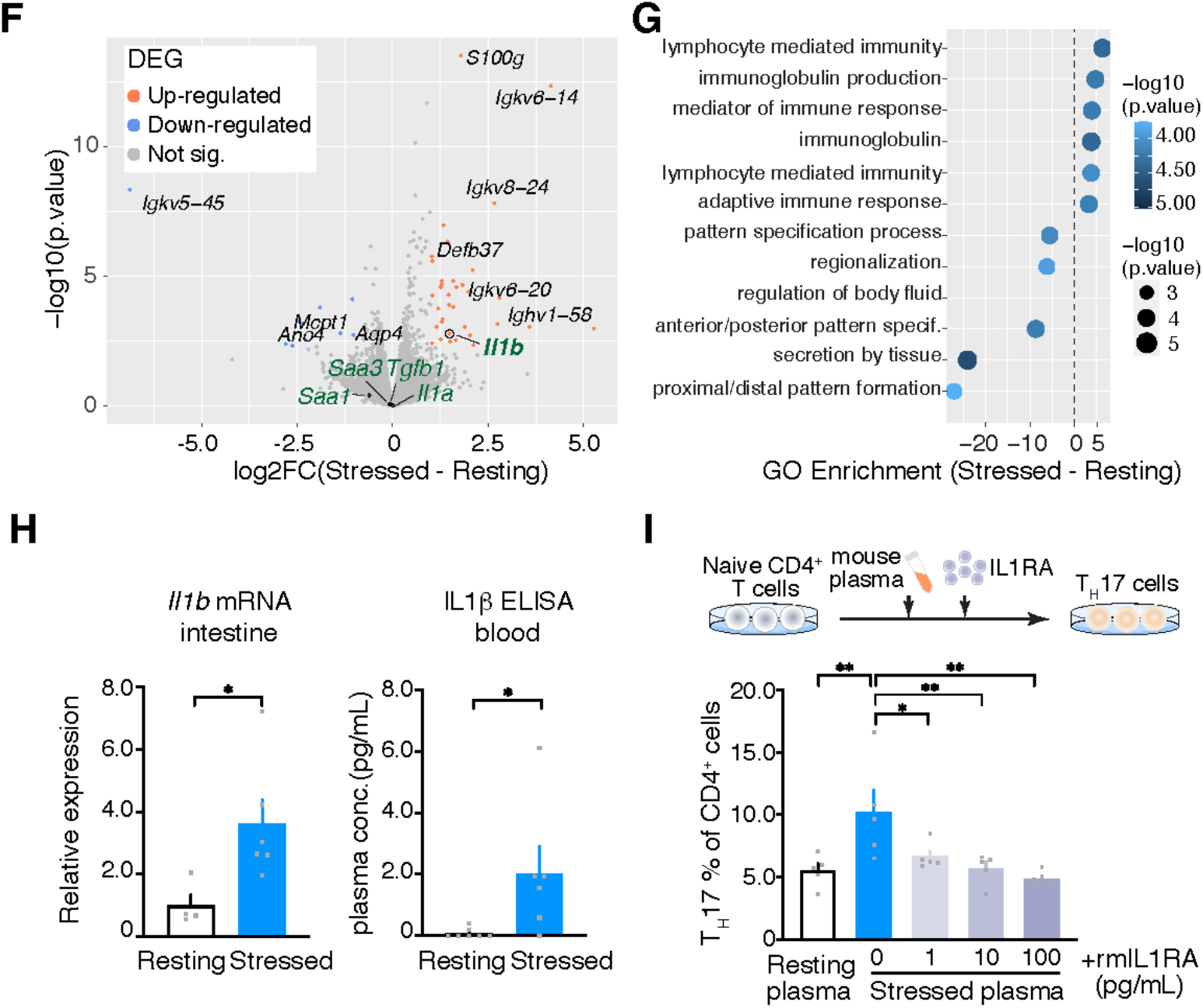
IL-22 is induced by stress via an IL-1β-T_H_17 pathway. **(A)** Intestinal permeability was determined by oral gavage of FITC-Dextran. n=6 for both groups. Microbial exposure was measured with stimulation of TLR4-SEAP reporter cells with plasma from resting and stressed mice. n=5 for both groups. **(B)** Circulating IL-22 concentrations before and after stress. Cytokines were profiled with the LegendPlex multiplex bead assay. n=8 for both groups. **(C)** Flow cytometry analysis of IL-22-producing T_H_17 cells in the intestine, showing representative flow plots (left) and quantification summary (right). n=5 for both groups. **(D)** *In vitro* differentiation of splenic naïve CD4^+^ T cells from resting and stressed mice into T_H_17 cells in the presence or absence of TH17-polarizing cytokines. n=5 for both groups. **(E)** *In vitro* differentiation of T_H_17 cells in the plasma of resting and stressed mice. n=5 for both groups. **(F)** Volcano plot showing differentially expressed genes in the intestines of resting and stressed mice. Detected T_H_17-skewing cytokines were highlighted in green. n=3 for each group. **(G)** Bubble plot showing top enriched GO pathways in the intestines of resting and stressed mice. n=3 for each group. **(H)** IL-1β levels in the intestine (mRNA) and blood (protein) before and after stress. n=4-6 for each group. **(I)** *In vitro* T_H_17 differentiation in the stressed mouse plasma with increasing concentrations of recombinant mouse IL1RA (rmIL1RA) to blocking of IL-1β. n=4 for all groups. Data are representative of 2-3 independent experiments and shown as Means ± SEMs. Statistical analysis was performed with Unpaired t-test (A-E, H) and 1-way ANOVA with Dunnett’s multiple comparisons (I). *P < 0.05; **P < 0.01; ***P < 0.001; ns, not significant.

To identify mechanisms by which stress induced T_H_17, we profiled the fecal microbiota between resting and stressed mice by 16S sequencing but failed to detect significant changes in the microbiota composition (bacteria and fungi) (Figure S4C and S4D), suggesting that the IL-22 and T_H_17 induced by repeated mild stress were unlikely due to microbial dysbiosis. We then isolated naïve CD4^+^ T cells from the spleens of stressed mice and examined their T_H_17 differentiation potential *in vitro* (Figure 4D). Naïve T cells from stressed mice were more prone to develop into T_H_17 cells than those from resting mice, consistent with increased T_H_17 cells in stressed mice. Since splenic T cells are exposed to circulating factors, we tested the effects of plasma from resting and stressed mice on T_H_17 cell differentiation. We found that plasma from stressed mice promoted naïve T cells from resting mice to develop into T_H_17 cells (Figure 4E). Because T_H_17 development is critically regulated by innate immune signals, we reasoned that stress-provoked gut leakage could lead to the production of T_H_17-skewing factors in the intestine and the blood.

To this end, we performed transcriptome analysis on the intestine tissues of resting and stressed mice. In agreement with stress-induced gut leakage, anti-microbial immune defense factors were significantly upregulated in the intestine after stress, such as S100, defensin and immunoglobin genes (Figure 4F, 4G and S4C). Importantly, the expression of IL-1β, a T_H_17-promoting cytokine downstream of microbial-sensing pathways, increased significantly in the intestine after stress (Figure 4F and 4H). No changes were detected for other known T_H_17-skewing factors including TGF-β1, serum amyloid As (SAAs), IL-6 and IL-23 in our setting (Figure 4F and S4C). Furthermore, the circulating IL-1β levels were also elevated after stress. To test whether IL-1β could account for the T_H_17-promoting effects of stressed mouse plasma, we antagonized IL-1β with its competitive inhibitor, IL1RA (Figure 4I). The addition of IL1RA to stressed mouse plasma blunted T_H_17 differentiation *in vitro* in a dose-dependent manner (Figure 4I). Taken together, our data suggested that stress promotes T_H_17 differentiation and IL-22 production through microbiota-stimulated IL-1β in the gut, thereby initiating a gut-brain pathway that protects mice from stress-induced anxiety disorders.

### IL-22 intervention reduces chronic stress-induced anxiety disorders in mice

Excessive chronic stress is a major etiological inducer of psychological diseases such as anxiety disorders and depression, which lack efficacious treatment and pose a huge challenge to modern societies. Thus, the IL-22-mediated immune-to-brain axis we identified here presents a promising therapeutic target for stress-induced psychological diseases. As our data thus far has shown an anxiolytic effect of IL-22 under a mild stress condition, we wanted to test whether IL-22 would confer protection against excessive chronic stress.

To this end, we subjected mice to an intense chronic stress paradigm (2-hour daily restraint for 21 consecutive days) known to induce mood disorders and intervened with daily injection of recombinant IL-22 (Figure 5A). In agreement with our previous data, IL-22 treatment led to a significant reduction (∼50%) of septal neuron activation (cFOS^+^) by stress (Figure 5A). While chronic stress led to anxiety disorders, IL-22 treatment significantly reduced the disease severity, suggesting IL-22 is also anxiolytic during chronic stress (Figure 5B). Furthermore, it has been reported that the antidepressants citalopram and mirtazapine can induce IL-22 from patient leukocytes *in vitro* (Munzer et al., 2013), suggesting that IL-22 might contribute to the anti-depressive benefits of these drugs. We then subjected IL-22-treated mice to the forced swimming test and tail suspension test, which are widely used to evaluate the therapeutic effects of antidepressants. We found that IL-22 treatment significantly reduced the immobility time of mice in both assays, suggesting that IL-22 ameliorated the development of depression induced by chronic stress (Figure 5C and 5D).

**Figure 5.**
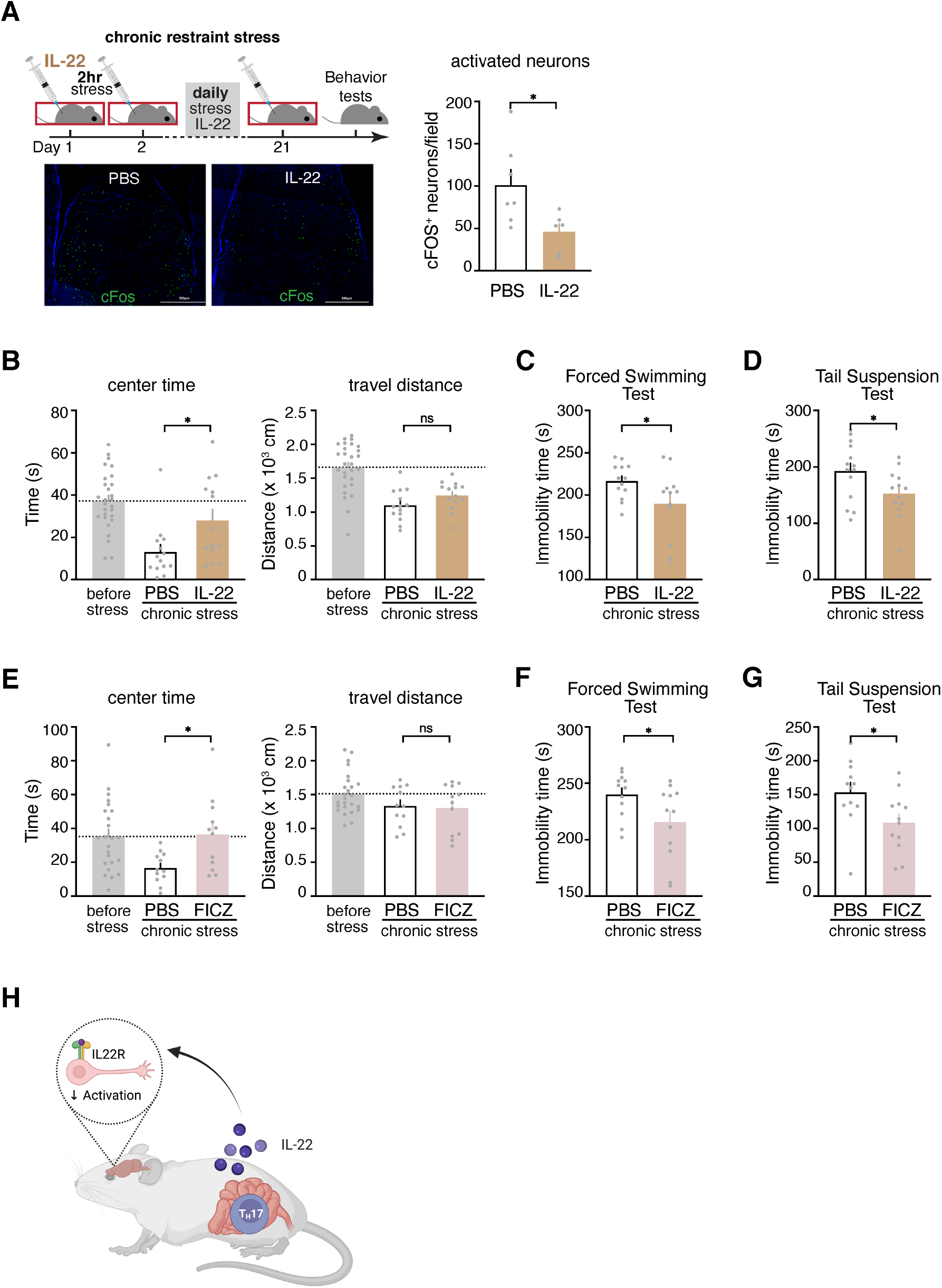
IL-22 protects mice from stress-induced psychological disorders. (**A-D**) Mice were subjected to excessive chronic stress, treated with IL-22, and analyzed for anxiety disorders and depression. (A) The experimental design and septal neuron activation in treated mice. Representative cFOS staining images (left) and quantification (right) are shown. n=7 and 6 for PBS and IL-22 treatment groups, respectively. (B) Anxiety was assessed with OFT. n=28, 14 and 14 for pre-stress, PBS and IL-22 treatment groups, respectively. (C,D) Depression was assessed with the forced swimming test (C) and tail suspension test (D). More immobility time indicates greater severity of depression in mice. n= 14 and 13 for the PBS and IL-22 treatment groups, respectively. (**E-G**) Mice were treated with the IL-22-inducing compound, FICZ, during excessive chronic stress and analyzed for anxiety disorders and depression. (E) Anxiety was examined with OFT. n=24, 12 and 12 for pre-stress, PBS and FICZ treatment groups, respectively. (F,G) Depression was assessed with the forced swimming test (F) and tail suspension test (G). n= 12 for both groups. (**H**) The working model. Data are presented as Means ± SEMs. Statistical analysis was performed with Unpaired t-test (A,C,D,F,G) and 1-way ANOVA with Dunnett’s multiple comparisons (B,E). *P < 0.05; ns, not significant.

Finally, because endogenous IL-22 can be modulated by diet-derived compounds (Gutierrez-Vazquez and Quintana, 2018), we tested whether the IL-22-inducing compound 6-formylindolo[3,2-b]carbazole (FICZ), a tryptophan derivative related to compounds enriched in cruciferous vegetables, could also increase stress resilience. As expected, treatment with FICZ during chronic stress significantly increased circulating IL-22 levels (Figure S5A). Similar to IL-22 treatment, FICZ treatment also reduced the development of anxiety disorders and depression induced by chronic stress (Figure 5E, 5F and 5G). Together, our data suggested that increasing IL-22 levels can protect animals against the accrued psychopathological effects of excessive chronic stress. Therefore, IL-22 intervention may provide a putative therapeutic strategy for stress-induced psychological diseases.

## Discussion

In summary, our work has revealed the cytokine IL-22 as the key mediator of an immune-to-brain axis defending against psychological stress. IL-22, and the T_H_17 cells that produce it, were induced by stress, following stress-provoked gut leakage and microbiota-stimulated IL-1β production. While IL-22 is well known to maintain gut barrier integrity, we have shown that it directly suppressed septal neurons that would otherwise drive anxiety-like behavior. We further demonstrated that IL-22 was both necessary and sufficient to reduce anxiety in mice in response to stress. In addition, manipulation of the IL-22 pathway by exogeneous IL-22 or a dietary compound ameliorated anxiety disorders and depression elicited by excessive chronic stress in mice. Indeed, the antidepressants citalopram and mirtazapine have been shown to induce IL-22 from human leukocytes *in vitro* (Munzer *et al*., 2013), suggesting that IL-22 might contribute to the therapeutic effects of these drugs. Therefore, our study suggests that the microbiota-stimulated IL-22 is critical for balanced allostatic responses to stress and represents an exciting opportunity to treat stress-related psychological diseases that have been increasingly prevalent in modern societies.

It has been a long-standing question regarding the physiological role of the immune system in the stress response. Mounting evidence suggests that pathological immune activation can lead to stress-related psychological diseases such as anxiety disorders and depression in animal models (Choi *et al*., 2016; Fan *et al*., 2019; Miyajima *et al*., 2017) and immune alterations have been observed in human patients with these diseases (Glaser and Kiecolt-Glaser, 2005; Menard *et al*., 2017). However, this may lie within the pathogenic extreme of the immune response spectrum and reflect the cumulative cost of immune activation during stress (Medzhitov, 2021). With our mild stress model of which the duration and intensity can be tolerated by WT mice, we observed initial immune activation and IL-1β expression in the gut following stress-provoked barrier dysfunction, which is known to increase microbiota exposure to the immune system (Shaler *et al*., 2021). IL-1β then induced T_H_17 differentiation and IL-22 production, which restored behavioral homeostasis in mice by suppressing septal neuron activation and anxiety. Therefore, the immune system senses microbiota danger signals during stress to regulate host neuronal and behavioral responses, thereby integrating environmental information to instruct host adaptation. Together, our study uncovers a previously-unappreciated defense role of the immune system against psychological stress. Note that T cells, the adaptive immune cells that take days to differentiate and function, might be optimally positioned to respond to chronic stress and defend against chronic stress-elicited psychological diseases. Further study of the immune-to-brain axis may yield new insights into how the immune system senses and responds to stressful encounters to ensure animal survival in its natural living environments.

## Acknowledgments

The authors thank Drs. Thomas S. Carroll and Ji-Dung Luo at the Bioinformatics Resource Center of The Rockefeller University for snRNA-seq data inspection and Dr. Christoph Kirst at UCSF for assistance with iDISCO data processing. The authors also thank other members of the Friedman laboratory and the Yu laboratory for discussions throughout this project.

## Funding sources

National Key Research and Development Program of China grant MOST-2021YFA0804703 (XY)

National Natural Science Foundation of China grant NSFC-32170928 (XY)

Howard Hughes Medical Institute Postdoctoral Fellowship of the Helen Hay Whitney Foundation (XY)

## Author contributions

Conceptualization: XY, JMF Methodology: AI, JL, LC, MS

Investigation: AI, MX, JL, LC, RS, CW, RF, ZJ, QQ, JL, BL, XY

Visualization: AI, JL, LC, ZJ, XY Funding acquisition: XY Supervision: XY

Writing – original draft: XY, AI, JMF

## Declaration of interests

The authors declare no competing interests.

## Notes

### Competing Interest Statement

The authors have declared no competing interest.

