## Supplemental data for "Microbiota-stimulated Interleukin-22 regulates brain neurons and protects against stress-induced anxiety"

### MATERIALS AND METHODS

#### Mice

C57BL/6J, *TCRb*$^{-/-}$ (B6.129P2-*Tcrb*$^{tm1Mom}$/J), *TRAP2* (STOCK *Fos*$^{tm2.1(icre/ERT2)Luo}$/J), Vgat-Cre (B6J.129S6(FVB)-*Slc32a1*$^{tm2(cre)Lowl}$/MwarJ) and *Il22ra1*$^{flox/flox}$ (B6.Cg-*Il22ra1*$^{tm1.1Koll}$/J) were obtained from the Jackson Laboratory (Bar Harbor, Maine) and maintained in Specific Pathogen Free (SPF) mouse facilities at Fudan University and/or The Rockefeller University. C57BL/6J and *TCRb*$^{-/-}$ mice were bred in-house after microbiota normalization by mixing dirty bedding. All mice were housed under 12hr:12hr light-dark (LD) cycles and fed *ad libitum*. All experimental procedures described in the study were approved by the Institutional Animal Care and Use Committees (IACUC) of Fudan University and/or The Rockefeller University.

#### Restraint stress

Male mice, aged 12-16 weeks, were individually housed for 1 week, followed by 2 minutes/day habituation to experimenter handling for 5 days. Mice were moved to a quiet procedure room and individually restrained in DecapiCones (Braintree Scientific, Braintree, MA) for 30 mins. Mice were then released back into their home cages and returned to the housing room. Three episodes of restraint were applied to mice on 3 consecutive days (1 episode/day). For extensive stress, mice were restrained for 2 hours per day and for a total of 21 days.

#### Open-field test (OFT)

The open-field arena (50 cm x 50 cm) was set up in a quiet, red light-lit room. At the beginning of experiments, individual mice were placed in a corner of the arena and recorded continuously by a high-definition video camera for 5 minutes before being returned to their home cages. The time of mouse spent in the center (23 cm x 23 cm) of the arena, total distance traveled, and velocity were analyzed with the EthoVision XT software (Noldus, Leesburg, VA) or the Tracking System software (Mobiledatum Shanghai, China).

**T cell reconstitution**

Spleens from wild-type C57BL/6J mice were harvested and macerated through a 70 μm cell strainer in ice-cold FACS buffer (DPBS, 3% FBS, 2 mM EDTA) to prepare single-cell suspension. Red blood cells were hypotonically lysed with 1x RBC lysis buffer (ThermoFisher, Waltham, MA). Total T cells were then purified with Dynabeads Untouched Mouse T Cells kit (ThermoFisher), following the manufacturer's instruction, with purity routinely over 95%. 2 x $10^6$ cells were transferred into each recipient mice by retro-orbital injection under anesthesia.

After recovering for 5 weeks, a small amount of blood was drawn from the tail veil and diluted into FACS buffer (1x DPBS, 3% FBS, 2mM EDTA). Samples were centrifuged at 300g for 5 minutes at 4 $^{o}$C to collect cells. Red blood cells were hypotonically lysed with the RBC lysis buffer (ThermoFisher). Cells were filtered through 40 μm cell strainers and spun down at 300g for 5 minutes. Cells were then blocked with anti-Mouse CD16/CD32 (2.4G2) for 30 minutes on ice and stained with CD45-FITC (30-F11), TCRβ-PE (H57-597), γδTCR-APC (GL3), CD4-PerCPCy5.5 (RM4-5) and CD8α-BV711 (53-6.7). Data were acquired with an LSR-II (BD Biosciences, Franklin Lakes, New Jersey) and Gallios (Beckman Coulter, Brea, CA) flow cytometers and analyzed with FlowJo v10.6.1 (BD) at the Flow Cytometry facilities of Fudan University and The Rockefeller University.

**LEGENDplex multiplex cytokine assay**

Blood was withdrawn from mice with Microvette® CB300 Capillary Blood Collection tubes (Sarstedt, 16.440.100) and centrifuged at 4000rpm for 20 minutes at 4 $^{o}$C. Plasma was taken into microcentrifuge tubes and stored at -80C until use. 25uL thawed plasma was assayed with the Mouse Th Cytokine LEGENDplex kit (Biolegend, 740005), following the manufacturer's instructions. Samples were analyzed with a BD LSR-II flow cytometer, demultiplexed with LEGENDplex Data Analysis Software Suite (Biolegend) and quantified with the drLumi package (Sanz et al., 2017) with the R and Bioconductor platform.

**iDISCO whole-mount immunolabeling and imaging**

Mice were stressed for 3 episodes of daily 30-minute restraint and sacrificed after the last treatment via transcardial perfusion with PBS and 4% paraformaldehyde (PFA) solution. Whole brains were dissected from the skulls and fixed in 4% PFA for another 24 hours. Clearing and immunolabeling were performed following the iDISCO+ protocol using an anti-cFos specific antibody (Synaptic Systems, 226003). Image acquisition was performed as described previously (Renier *et al.*, 2014) using the LaVision Ultramicroscope II, with cFos labeling acquired in the 640 nm channel and background autofluorescence acquired in the 480 nm channel. Images were reconstructed, mapped to the Allen Brain Reference Atlas, and quantified with the ClearMap pipeline (https://www.idisco.info/). Z scores were calculated and mapped to a tree object depicting the Allen brain atlas hierarchy using R packages "igraph" and "ggraph". Heatmap was generated with z scores of respective regions with the R package "gplots".

**RNA-seq of the septum**

RNA-seq samples were prepared as published before (Knight et al., 2012), with modifications. Briefly, mice were treated with 30-minute restraint stress for 3 days and sacrificed 1.5 hours after the last restraint. Brains were dissected and immediately immersed into ice-cold Dissection Buffer (1x HBSS, 2.5 mM HEPES pH7.4, 4 mM $NaHCO_3$ and 35 mM Glucose). Tissues from 4-5 mice were pooled and transferred into Teflon-glass homogenizer with Homogenization Buffer [10 mM HEPES pH7.4, 150 mM KCl, 5 mM $MgCl_2$, 0.5 mM DTT, 0.5

U/mL SUPERase•In (ThermoFisher, AM2696), 0.5 U/mL RNase Inhibitor (ThermoFisher, AM2682) and 2x EDTA-free Halt protease inhibitor cocktail (ThermoFisher, 87786)]. Tissues were then dounced 12 strokes or until no visible clumps of tissue could be seen. The lysate was then transferred into a new RNase-free microcentrifuge tube and cleared by centrifuging at 2000g at 4 °C for 10 minutes. Clear supernatant was then supplemented with 0.6% NP-40 and 18 mM DHPC and incubated on ice for 2 minutes. Samples were centrifuged again at 16,000g at 4C for 10 minutes and clear supernatant was collected into new microcentrifuge tubes. RNA was extracted with an RNeasy Micro Kit (Qiagen, 74004) RNA quality was determined by a Bioanalyzer 2000 (Agilent) and samples with RIN>9 were used to prepare sequencing libraries.

Raw RNA-seq data were aligned to the mm10 mouse reference genome with HISAT2 to generate gene expression matrices. Differentially expressed genes were identified with the R package “Limma-Voom” with paired analysis. Pathway analysis was performed with the Gene Set Enrichment Analysis (GSEA) software (Broad Institute), coupled with the curated mouse pathway database MousePath (downloaded from http://ge-lab.org/gskb/). Expression data of Azevedo *et al* (Azevedo *et al.*, 2020) was downloaded from Gene Expression Omnibus (GSE154749) and analyzed with “Limma-Voom”. Stressed-induced genes were identified as *p.value*<=0.01 and *log2FC*>=0.67 and formatted into an MSigDB signature for GSEA analysis.

**Stereotactic surgery and cannula implantation**

Brain regions were identified according to the Allen mouse brain atlas and coordinates were relative to the bregma. Mice were anesthetized with 3.5% isoflurane and switched to 2% isoflurane during the surgery. Mice were mounted onto a KOPF 942 stereotaxic frame (Kopf Instruments, Tujunga, CA) with heads leveled. Skulls were exposed by incising the scalp with a sterile scalpel and cleaned with 30% $H_2O_2$. A small hole was drilled into the skull over the septum (AP=0.58 mm, ML=0.00 mm). For AAV injection, 0.5-1 µL virus was injected at a rate of 0.1-0.2 µL/min with a Hamilton syringe (DV=3.50 mm), and the syringe was held in place for another 3 min to allow sufficient viral diffusion. For cannula implantation, guide cannulas (P1 Technologies, Roanoke, VA) were inserted with the same coordinates as above and secured to the skull with Metabond dental cement (Parkell Inc, Edgewood, NY). Dummy cannulas were installed to prevent dust. The skin was closed with glue. All surgical procedures were performed under aseptic conditions and mice were monitored up to 72 hrs post-surgery to ensure full recovery. Behavior analysis was carried out at least 2 weeks later.

**IL-22 and FICZ injection**

Recombinant mouse IL-22 (Biolegend 576202 and Novoprotein C047) was reconstituted in sterile PBS, aliquoted, and stored at -80 °C. For microinjection into the septum, the IL-22 stock was diluted to 4 ng/µL with saline and 0.25 µL (1 ng) was injected at 0.125 µL/min with a Harvard syringe pump and a 1 uL Hamilton syringe connected to an internal cannula. Mice were immobilized for another 5 minutes before the internal cannula was taken out to avoid backflow. For intraperitoneal (i.p.) injection, the IL-22 stock was diluted to 10 µg/mL with PBS and 100 µL (2 µg) was injected for each mouse. FICZ was i.p. injected at 1 ug/mouse every 3 days or 4 days.

**TRAP2 labeling and reactivation of septal neurons**

The AAV5-hSyn-DIO-hM3D(Gq)-mCherry (Addgene, 44361) virus was stereotactically injected into the septum of TRAP2 mice as described above. Mice were given at least 2 weeks to recover and habituated to intraperitoneal injection by daily injection of 200 µL PBS for 5 days. Mice were injected with 20mg/kg of 4-hydroxytamoxifen (4-OHT) and randomly assigned into 2 groups, with one group concurrently restrained for 30 minutes in DecapiCones (the Stress labeling group) and the other group returned to their home cages immediately (the Resting labeling group). The virus-encoded hM3D(Gq) channel was allowed to express for 3 weeks before behavioral analysis. For individual mice, saline was first injected to determine basal anxiety in the OFT, followed by injection of clozapine N-oxide (CNO) at the dose of 1 mg/kg to determine neuronal reactivation-elicited anxiety. Changes in the time that individual mice spent in the center of the OFT arena with saline and CNO injection were calculated to evaluate the effects of septal neuron reactivation.

**Acute-brain slice preparation and patch-clamp recordings**

Mice were deeply anesthetized using pentobarbital sodium (50 mg/kg) before decapitation and rapid removal of the entire brain into an ice-cold, oxygenated (95% O2/5% CO2) cutting solution (220 mM sucrose, 3 mM KCl, 5 mM $MgCl_2$, 1 mM $CaCl_2$, 1.25 mM $NaH_2PO_4$, 26 mM $NaHCO_3$, 10 mM glucose [pH 7.30], 310–320 mOsm/l). Coronal slices containing the affected brain area were cut at 300 μm using a vibrating microtome (Dosaka, Kyoto, Japan) and incubated in oxygenated (95% $O_2$/5% $CO_2$) artificial cerebrospinal fluid (aCSF) (125 mM NaCl, 2.50 mM KCl, 2.50 mM $CaCl_2$, 1.50 mM $MgSO_4$, 1 mM $NaH_2PO_4$, 26.20 mM $NaHCO_3$, 11 mM glucose [pH 7.30], 300–310 mOsm/l) for 1 h at 32°C and then stored at room temperature.

Recordings of slices were carried out at room temperature. Neurons were selected for electrophysiological recording located in the lateral septum. Slices were kept fully submerged during recording and continuously perfused (3–4 ml/min) with 95% $O_2$/5% $CO_2$-equilibrated ACSF. Recording electrodes were pulled from borosilicate glass on a P-97 four-stage puller (Sutter Instruments, Novato, CA, USA), with a resistance of 3–6 MΩ when filled with an internal solution. Whole-cell patch-clamp recordings were obtained under visual guidance by differential interference microscopy (Zeiss, Examiner.A1).

Miniature inhibitory postsynaptic currents (mIPSCs) were recorded using voltage-clamp mode in ACSF with 25 mM DL-2-Amino-5-phosphonopentanoic acid (DL-AP5) (Sigma, USA),12.5 μM CNQX (Sigma, USA) and 1 μM TTX (Sigma, USA). Recording pipettes were filled with an internal solution containing 125 CsCl, 8 NaCl, 0.6 ethylene glycol tetraacetic acid (EGTA), 10 4-(2-hydroxyethyl)-1-piperazineethanesulfonic acid (HEPES), 4 Mg-ATP, 0.3 $Na_3$-GTP, 10 Na-phosphocreatine, and 2 QX-314 (Abcam, England) (pH 7.30, 290 mOsm/l). mIPSCs were analyzed using Mini-Analysis software (Synaptosoft, Decatur, Georgia, USA). Resting potential and action potential were recorded using the current-clamp mode in ACSF, and recording pipettes were filled with an internal solution containing 150 mM K-gluconate, 0.40 mM EGTA, 10 mM HEPES, 2 mM Mg-ATP, 0.10 mM $Na_2$-GTP, and 8 mM NaCl, (pH 7.30, 290 mOsm/l). 100 pA injection currents of 1000 ms duration were used to induce action potentials. Action potential analysis was carried out using pClamp 10.7 (Axon Instruments). Signals were amplified and filtered (2 kHz) using an Axopatch 700B amplifier, sampled at 10 kHz using a Digidata 1550B, and recorded with Clampex (Molecular Devices). Only cells with a stable access resistance of < 30 MΩ throughout the recording period were included in the analysis.

**Single-nucleus RNA-Seq (snRNA-seq)**

Nuclei were prepared as reported before (Krishnaswami et al., 2016) with modifications. Specifically, septal tissues were dissected from 10 mice, pooled together, and stored in 1mL of ice-cold Brain Homogenization Buffer [10 mM Tricine-KOH pH8, 25 mM KCl, 5mM $MgCl_2$, 250 mM Sucrose and 0.1% Triton X-100), supplemented with 0.5 U/mL SUPERase•In (ThermoFisher, AM2696), 0.5 U/mL RNase Inhibitor (ThermoFisher, AM2682) and 2x EDTA-free Halt protease inhibitor cocktail (ThermoFisher, 87786)]. Samples were transferred into a cold Wheaton 1mL glass homogenizer and ground by 10 strokes with the "Loose" pestle, followed by 14 strokes with the "Tight" pestle. Homogenate was then filtered with 40um cell strainers and underlaid with 1mL 20% iodixanol in a clear ultracentrifuge tube. The sample was centrifuged at 10,000 g for 20 min at 4 °C with a low acceleration/deceleration setting. Nuclei was resuspended in 500uL Staining Buffer (1x DPBS, 0.5% BSA, 0.1% Triton X-100, 0.05% NP-40, 0.5 U/mL SUPERase•In, 0.5 U/mL RNase Inhibitor, 2x EDTA-free Halt protease inhibitor cocktail) and stained with anti-NeuN-AF647 (Abcam, ab190565, 1:500) and Hoechst 33342 (ThermoFisher, H3570, 1:500) at 4 °C for 30 minutes. Nuclei were washed once with Staining Buffer, spun down at 1000g for 5 minutes and resuspended in Nuclei FACS Buffer (1x DPBS, 0.5% BSA, 0.5U/mL SUPERase•In, 0.5U/mL RNase Inhibitor, 2x EDTA-free Halt protease inhibitor cocktail). NeuN$^+$ nuclei were collected with a BD FACSAria cell sorter, inspected with a Countess automated cell counter, and loaded into a 10x Genomics Chromium controller. Libraries were prepared by the Genomics Resource Center of The Rockefeller University with a Chromium Single Cell 3' kit v2 (10x Genomics) according to the manufacturer's instructions. Initial data inspection was performed by the Bioinformatics Resource Center (BRC) at The Rockefeller University.

**snRNA-seq data analysis**

Single nucleus sequencing was performed using the Chromium platform from 10X Genomics, and the reads were generated on an Illumina Nextseq sequencer. A total of 349 million raw read pairs were generated. For analysis, reads were first processed by cellranger from the 10X Genomics website (https://support.10xgenomics.com/), following default parameters. In particular, an intron-to-exon converted reference was used for single nucleus data. Then the cellranger generated matrix file was loaded into Seurat v3 (Stuart et al., 2019).

Downstream analysis generally followed the recommended Seurat protocols. In brief, the data is first filtered to contain only cells with 500 to 6000 genes per cell and no more than 2% of mitochondria mRNA per cell. It was then log-normalized and scaled and the top 2000 highly variable genes were identified. Dimension reductions were performed on these highly variable genes with the first PCA then UMAP. Then the cells were clustered and could be visualized in various plots. Due to the high drop-out nature of single-cell/nucleus expression data, imputation of gene expression was performed using MAGIC (van Dijk et al., 2018).

**Intestinal immune cell analysis**

Intestinal immune cells were isolated and analyzed as previously described (Yu et al., 2013). For cytokine analysis, single-cell suspensions were stimulated with 50 ng/mL phorbol12-myristate-13-acetate (PMA, Sigma-Aldrich, P8139) and 500 ng/mL ionomycin (Sigma-Aldrich, I0634) in the presence of 1 μg/mL brefeldin A (BFA, Sigma-Aldrich, B6542) for 4 hours at 37C. Cells were then stained with Zombie NIR (Biolegend) for 10 minutes at 4 °C and blocked with anti-Mouse CD16/CD32 (2.4G2) for 30 minutes on ice. Cells were then processed with the BD Cytofix/Cytoperm Kit and stained with IL17A-eFluor450 (17B7), CD45-BV570 (30-F11), CD3ε-BV650 (500A2), CD8α-BV711 (53-6.7), CD90.2-BV785 (30-H12), CD4-FITC (RM4-5), γδTCR-PerCP-eFluor710 (GL3), IL22-PE (1H8PWSR), TCRβ-PE-TexasRed (H57-597), IFNγ-PECy7 (XMG1.2), CD127-APC (A7R34), TNFα-AlexaFluor700 (MP6-XT22) and NK1.1-APC-eFluor780 (PK136). For transcription factor staining, cells were processed with the Thermo Fisher Transcription Factor Fix/Perm Kit and stained with RORγt-BV421 (Q31-378), CD4-BV510 (RM4-5), CD45-BV570 (30-F11), CD11b-BV605 (M1/70), CD3ε-BV650 (500A2), CD8α-BV711 (53-6.7), CD90.2-BV785 (30-H12), Foxp3-FITC (FJK-16s), T-bet-PE (4B10), CD19-PE-Cy5 (6D5), FcεR1-PE-Cy7 (MAR-1), CD11c-APC (HL3), GATA3-eFluor660

(TWAJ), TCRβ-AlexaFluor700 (H57-597) and NK1.1-APC-eFluor780 (PK136). Data were acquired with an Aurora flow cytometer (Cytek, Bethesda, MD) and analyzed with FlowJo v10.6.1 (BD).

***In vitro* $\mathbf{T_H}$17 polarization and analysis**

Naïve mouse $CD4^+$ T cells were isolated with the MojoSort™ Mouse CD4 Naïve T Cell Isolation Kit (Biolegend, 480040) following the manufacturer's instructions. *In vitro* $T_H$17 polarization was performed as described before with minor modifications (Yu *et al.*, 2013). Briefly, a 96-well plate was coated with 10 μg/mL of anti-Hamster IgG antibody (Jackson Immunoresearch, 127-005-099) at 4 °C overnight. 100,000 naïve mouse $CD4^+$ T cells were seeded into each well in X-VIVO 15 media (Lonza, BEBP02-061Q), supplemented with 0.25 μg/mL anti-mouse CD3ε (145-2C11, Biolegend, 100359), 1 μg/mL anti-mouse CD28 (37.51, Biolegend, 102121), 2.5 μg/mL anti-mouse IL-4 (11B11, Biolegend, 504101), 2.5 μg/mL anti-mouse IFNγ (XMG1.2, Biolegend, 505833) and 5% (vol/vol) plasma harvested from control and stressed mice with heparinized capillary tubes. For $T_H$17polarization, 20 U/mL of recombinant mouse IL-2 (Biolegend, 575402), 20 ng/mL recombinant mouse IL-6 (Biolegend, 575702) and 0.3 ng/mL recombinant mouse TGF-β (Biolegend, 575702) were added into the culture media. For IL-1β blocking experiments, recombinant mouse IL1RA was supplemented in the culture media at the concentrations of 1, 10 and 100 pg/mL. Cells were incubated at 37 °C for 72 hours and stimulated with PMA, Ionomycin, and BFA for 4 hours. Cells were stained with CD4-eFluor506 (RM4-5), RORγt-BV421 (Q31-378), and IL17A-APC (17B7) with the True-Nuclear Transcription Factor Buffer (Biolegend, 424401) or eBiosciences Foxp3/Transcription Factor Staining kit (ThermoFisher, 00-5523-00). $T_H$17 cells were analyzed with a Gallios flow cytometer (Beckman Coulter) and quantified with FlowJo v10.6.1 (BD Biosciences).

**Enzyme-Linked Immunosorbent Assay (ELISA)**

Plasma was collected as described above and ELISA was performed following the manufacturers' instructions. The following ELISA kits were used: ELISA MAX™ Mouse IL-22 Deluxe Set (Biolegend, 436304), IL-1 beta Mouse Uncoated ELISA Kit (ThermoFisher, 88-7013-88).

**Reverse transcription-polymerase chain reaction (RT-PCR)**

Mouse intestinal tissues were dissected and immediately frozen in liquid nitrogen. Tissues were homogenized with a Wiggens D-130 Handheld Homogenizer and RNA was purified with an RNAsimple Total RNA Isolation Kit (Tiangen, Q711-03). Total RNA was then reversely transcribed with a HiScript III 1st Strand cDNA Synthesis Kit (+gDNA wiper) (Vazyme, R312-01). Quantitative PCR was performed with gene-specific primers (Table S1) and ChamQ Universal SYBR qPCR Master Mix (Vazyme, Q711-03) in a QuantStudio 7 Flex Real-Time PCR machine (ThermoFisher). Gene expression was calculated as $-2^{\Delta Ct}$ with *Gapdh* as the internal control.

**Immunohistochemistry (IHC) staining**

Mice were transcardially perfused with PBS for 5 minutes and 4% PFA for another 5 minutes. Brains were dissected, fixed in 4% PFA overnight at 4 °C, and sliced with a VT1000S vibrating blade microtome (Leica). Free-floating sections (50 μm thick) were permeabilized and blocked with IHC staining solution (PBS, 3% BSA, 2% normal goat serum, 0.3% Triton X-100) for 2 hours at room temperature and stained with an anti-cFOS antibody (clone# 9F6, 1:500, Cell Signaling, 2250) for 48 hours at 4 °C with gentle shaking. Sections were then washed 3 times with IHC washing solution (PBS, 0.1% Triton X-100) for 10 minutes and stained with a goat anti-rabbit IgG-AlexaFluor488 secondary antibody (1:1000, ThermoFisher, A11008) for 1 hour at room temperature. Sections were washed again, mounted in DAPI Fluoromount-G (YEASEN, 36308ES11), and imaged with a Nikon A1 confocal microscope. Images were then processed with the Fiji ImageJ software (NIH).

**Intestinal permeability assessment**

Mice were restrained for 30 min/day for the first 2 days. On day 3, mice were orally gavaged with 600 mg/kg Dextran-FITC and restrained for 30 min. Blood was collected through retro-orbital bleeding 4 hrs later and centrifuged at 4000rpm for 20 minutes at 4 $^{o}$C. FITC intensities in the plasma were measured at 528 nm emission with a Biotek Synergy 2 Multi-Mode Plate Reader. In other experiments, plasma from resting and stressed mice were added to the TLR4-SEAP reporter (HEK-Blue™-4, Invivogen, rep-lps2) cells at 10% v/v and incubated for 16 hrs. SEAP activities were determined with the H5N1Alkaline Phosphatase Assay Kit (Beyotime, P0321M) following the manufacturer's instructions.

**Statistical analysis**

Statistical analysis was performed with GraphPad Prism, R, and Python with packages specified in individual experiments. All data are presented as Mean ± SEM unless otherwise noted. Significance is defined as *P<0.05, **P<0.01 and ***P<0.001.

**Data and materials availability**

Septal tissue RNA-seq and snRNA-seq data will be submitted to GEO under access GSE193863 and GSE193890, respectively. Mouse strains are all commercially available. All other data are available in the main text or supplementary materials.

**Supplemental information**

Figures S1 to S5

Tables S1

References

1 Figure S1

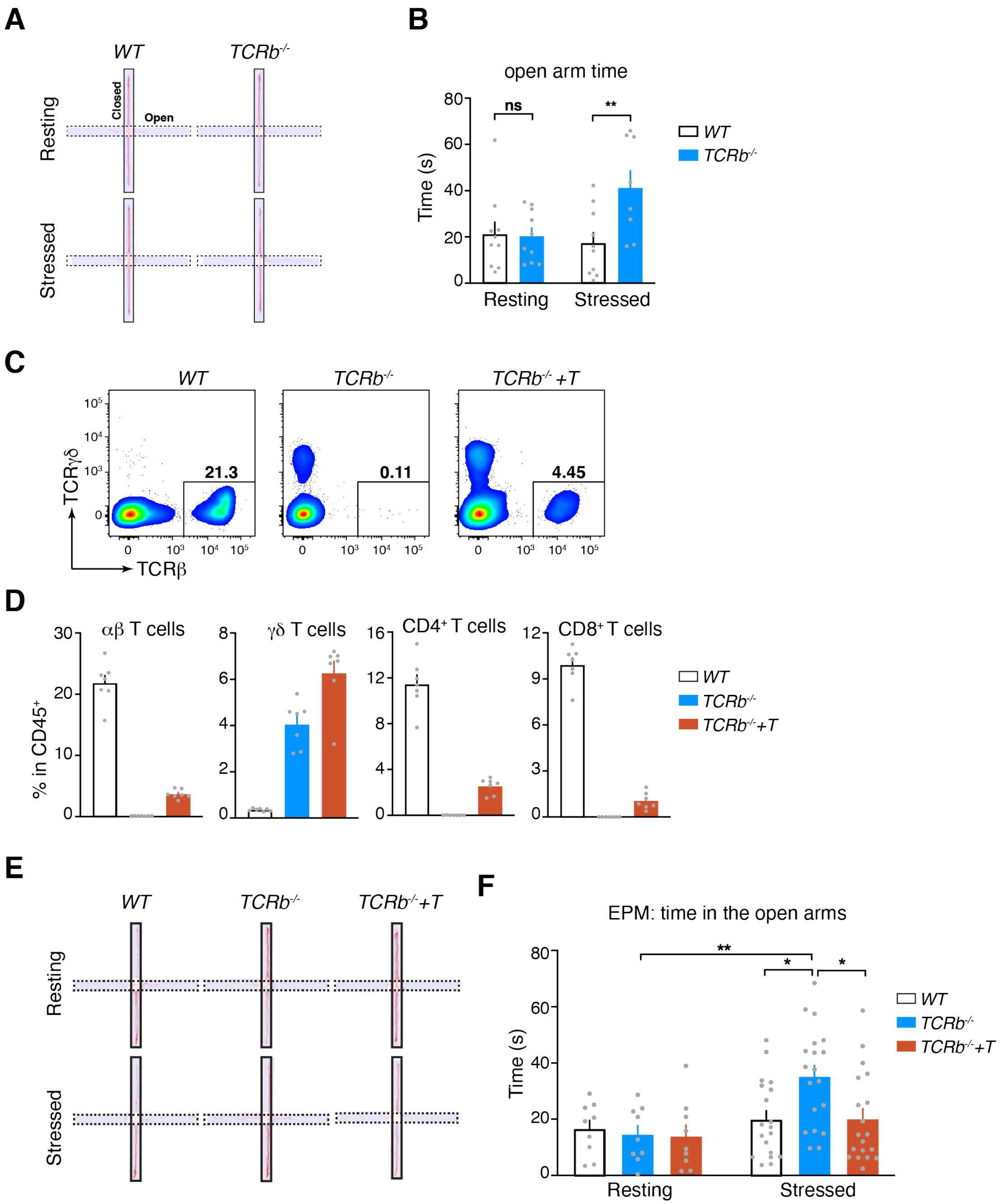
A
WT
TCRb-/-
Resting
Stressed
Closed
Open
B
open arm time
Time (s)
ns
**
Resting
Stressed
WT
TCRb-/-
C
WT
TCRb-/-
TCRb-/- +T
21.3
0.11
4.45
TCRγδ
TCRβ
D
αβ T cells
γδ T cells
CD4+ T cells
CD8+ T cells
% in CD45+
WT
TCRb-/-
TCRb-/-+T
E
WT
TCRb-/-
TCRb-/-+T
Resting
Stressed
F
EPM: time in the open arms
Time (s)
Resting
Stressed
WT
TCRb-/-
TCRb-/-+T

**Figure S1. T cell deficiency in *Tcrb*$^{-/-}$ mice reduces stress-provoked anxiety.**

(**A,B**) WT and *Tcrb*$^{-/-}$ mice were subjected to repeated mild restraint stress (30-minutes restraint/day for 3 days) and anxiety-like behavior was assessed with Elevated Plus Maze (EPM). (A) Representative track plots of mice in EPM. (B) Quantification of time that mice spent in the open arms of EPM, which is indicative of anxiety levels. n=10, 10, 10, 8 for WT-resting, *Tcrb*$^{-/-}$-resting, WT-stressed and *Tcrb*$^{-/-}$-stressed, respectively.

(**C,D**) *Tcrb*$^{-/-}$ mice were reconstituted with WT splenic T cells. (C) Representative flow plots of mouse blood. (D) Frequencies of αβ-, γδ-, CD4$^+$ and CD8$^+$ T cells in the blood. T cell transfer partially rescued T cell deficiency in *Tcrb*$^{-/-}$ mice. n=7 for all 3 groups.

(**E,F**) WT, *Tcrb*$^{-/-}$ and T cell-rescued *Tcrb*$^{-/-}$ (*Tcrb*$^{-/-}$+*T*) mice were treated with restraint stress and analyzed in EPM. (E) Representative track plots of mice in EPM. (F) Quantification of time that mice spent in the open arms of EPM. n=9 for resting groups and 19 for stressed groups.

Data are presented as Means ± SEMs. Statistical analysis was performed with 2-way ANOVA with Sidak's multiple comparisons (B,F). $*P < 0.05$; $**P < 0.01$; ns, not significant.

### Figure S2

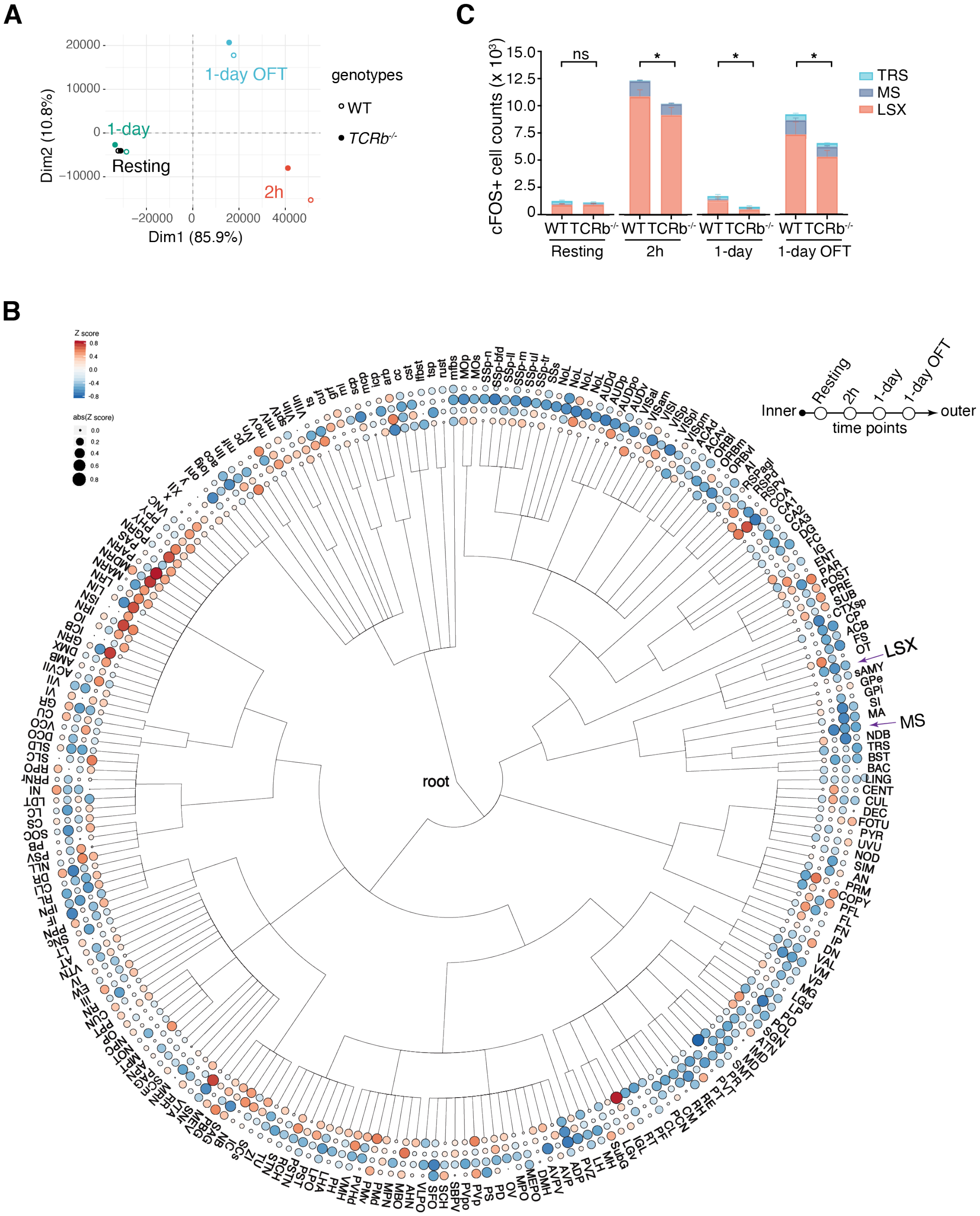
A
Dim1 (85.9%)
Dim2 (10.8%)
1-day OFT
1-day
Resting
2h
genotypes
WT
TCRb-/-
B
Z score
abs(Z score)
root
LSX
MS
Inner
Resting
2h
1-day
1-day OFT
outer
time points
C
cFOS+ cell counts (x 10³)
ns
*
TRS
MS
LSX
WT
TCRb-/-
Resting
2h
1-day
1-day OFT

### Figure S2 (cont'd)

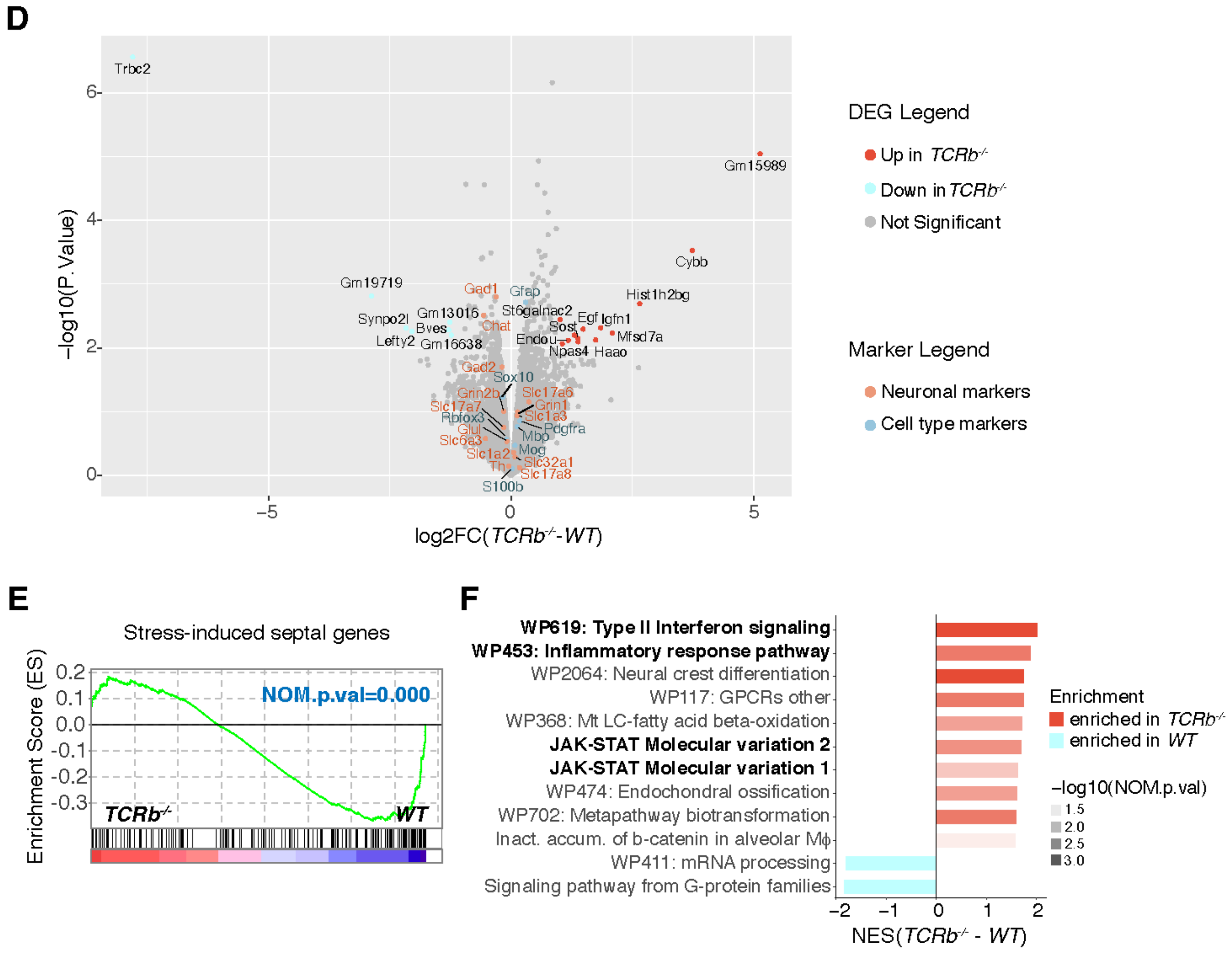

**Figure S2. The septal area is differentially activated by stress in WT and *Tcrb$^{-/-}$* mice.**

(**A-C**) Neuronal activation was assessed by whole-brain imaging of cFOS expression with iDISCO. WT and *Tcrb$^{-/-}$* mouse brains were collected at baseline (Resting), two hours after the final restraint stress (2h), one day after the final restraint stress (1-day), and after exposure to OFT one day after the final restraint stress (1-day OFT). n=5 for each group. (A) Principal component analysis (PCA) of mean cFOS$^+$ cell counts of brain regions for WT and *Tcrb$^{-/-}$* mice at all 4 time points. (B) Tree plot showing differential activation of brain regions by stress in WT and *Tcrb$^{-/-}$* mice (z scores) at each time point. Blue dots indicate negative z scores and reduced neuron activation in *Tcrb$^{-/-}$* mice and red dots indicate positive z scores and increased neuron activation in *Tcrb$^{-/-}$* mice. The intensity of colors and size of dots reflects the absolute values of the z score. Dots from the inner circle to the outer circle correspond to z scores of the Resting, 2h, 1-day and 1-day OFT timepoints. (C) cFOS$^+$ cell counts in the septal area of WT and *Tcrb$^{-/-}$* mice, including the lateral septum (LSX), medial septum (MS), and triangular septal nucleus (TRS).

(**D-F**) RNA-seq analysis of septal tissues in stressed WT and *Tcrb$^{-/-}$* mice. n=2 for each genotype. (D) Volcano plot showing differential septal gene expression between WT and *Tcrb$^{-/-}$* mice, with known cell- and neuron-type markers highlighted. (E) GSEA analysis of enrichment of a stress-induced gene signature derived from Azevedo *et al* in WT and *Tcrb$^{-/-}$* mice. (F) GSEA analysis of signaling pathways enriched or depleted in *Tcrb$^{-/-}$* mice.

Data are presented as Means ± SEMs. Statistical analysis was performed with 2-way ANOVA with Sidak's multiple comparisons (C). *$P < 0.05$; ns, not significant.

### Figure S3

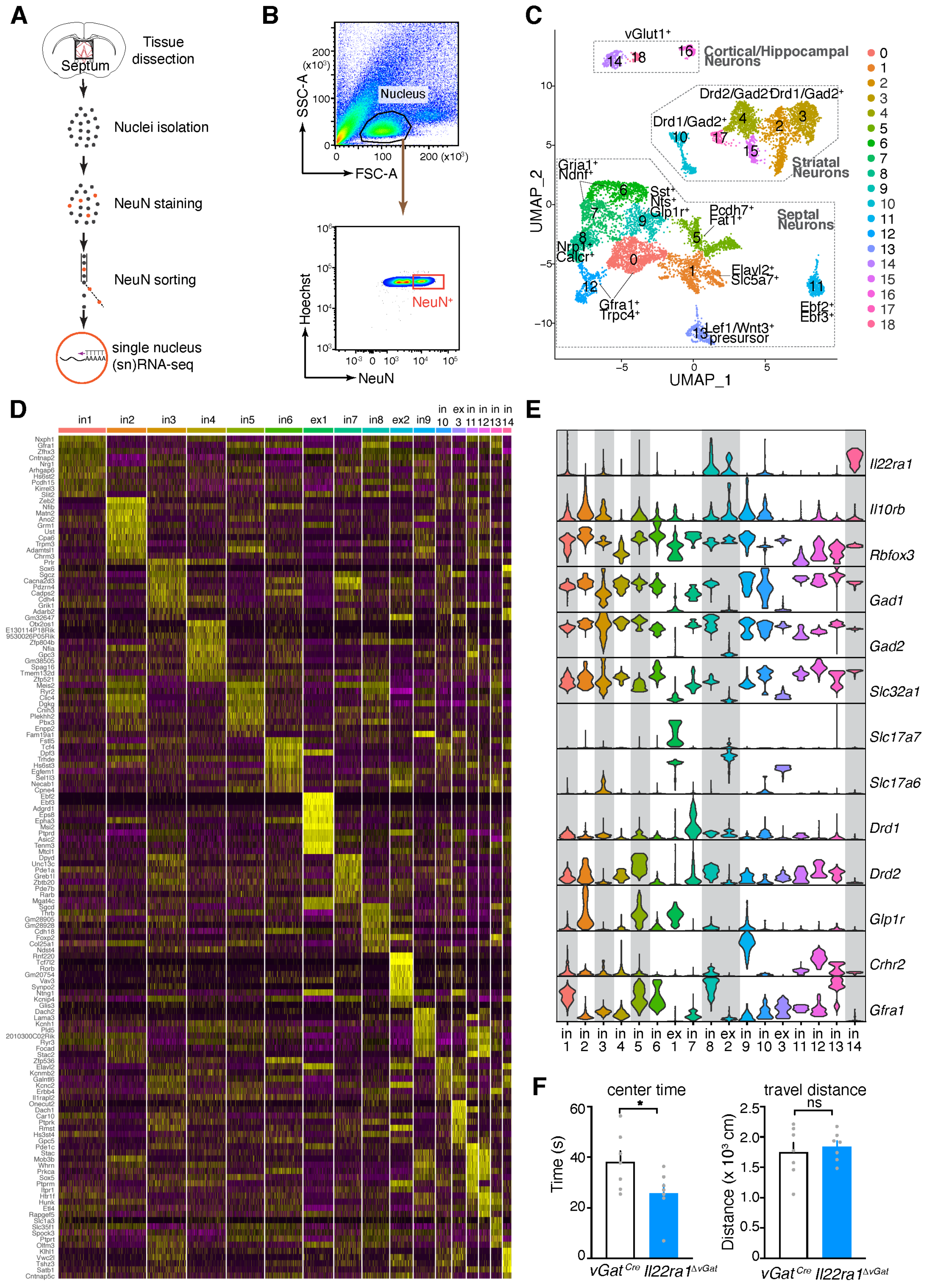
A
Tissue dissection
Septum
Nuclei isolation
NeuN staining
NeuN sorting
single nucleus (sn)RNA-seq
B
SSC-A
FSC-A
Nucleus
Hoechst
NeuN
NeuN+
C
vGlut1+
Cortical/Hippocampal Neurons
Drd2/Gad2+
Drd1/Gad2+
Striatal Neurons
Grja1+ Ndnf+
Sst+ Nts+ Glp1r+
Pcdh7+ Fat1+
Septal Neurons
Nrp1+ Calcr+
Elavl2+ Slc5a7+
Gfra1+ Trpc4+
Ebf2+ Ebf3+
Lef1/Wnt3+ presursor
UMAP_1
UMAP_2
D
in1 in2 in3 in4 in5 in6 ex1 in7 in8 ex2 in9 in 10 ex 3 in 11 in 12 in 13 in 14
E
Il22ra1
Il10rb
Rbfox3
Gad1
Gad2
Slc32a1
Slc17a7
Slc17a6
Drd1
Drd2
Glp1r
Crhr2
Gfra1
F
center time
Time (s)
travel distance
Distance (x 10³ cm)
ns
vGat Cre
Il22ra1 ΔvGat

**Figure S3. Septal neurons express IL-22 receptors.**

(**A-E**) snRNA-seq analysis of septal neurons. (A) Illustration of the snRNA-seq procedure. (B) Representative flow plots of nuclei sorting. (C) Cell clusters visualized by Uniform Manifold Approximation and Projection (UMAP) embedding. Neurons from the septum and neighboring areas are demarcated by dashed lines. (D) Heatmap showing signature genes of septal clusters that were identified by reanalysis of septal neurons. (E) Violin plots showing the expression of IL-22 receptor subunits (*Il22ra1* and *Il10rb*) and neuron marker genes. Clusters with both *Il22ra1* and *Il10rb* expressed in more than 5% of member cells are shaded in gray.

(**F**) Mice with *Il22ra1*-deficiency in $vGat^{+}$ neurons were assayed for anxiety-like behavior in OFT. Center time and travel distance are quantified. n= 7 for both groups.

Data are presented as Means ± SEMs. Statistical analysis was performed with the Unpaired t-test (F). *P < 0.05; ns, not significant.

### Figure S4

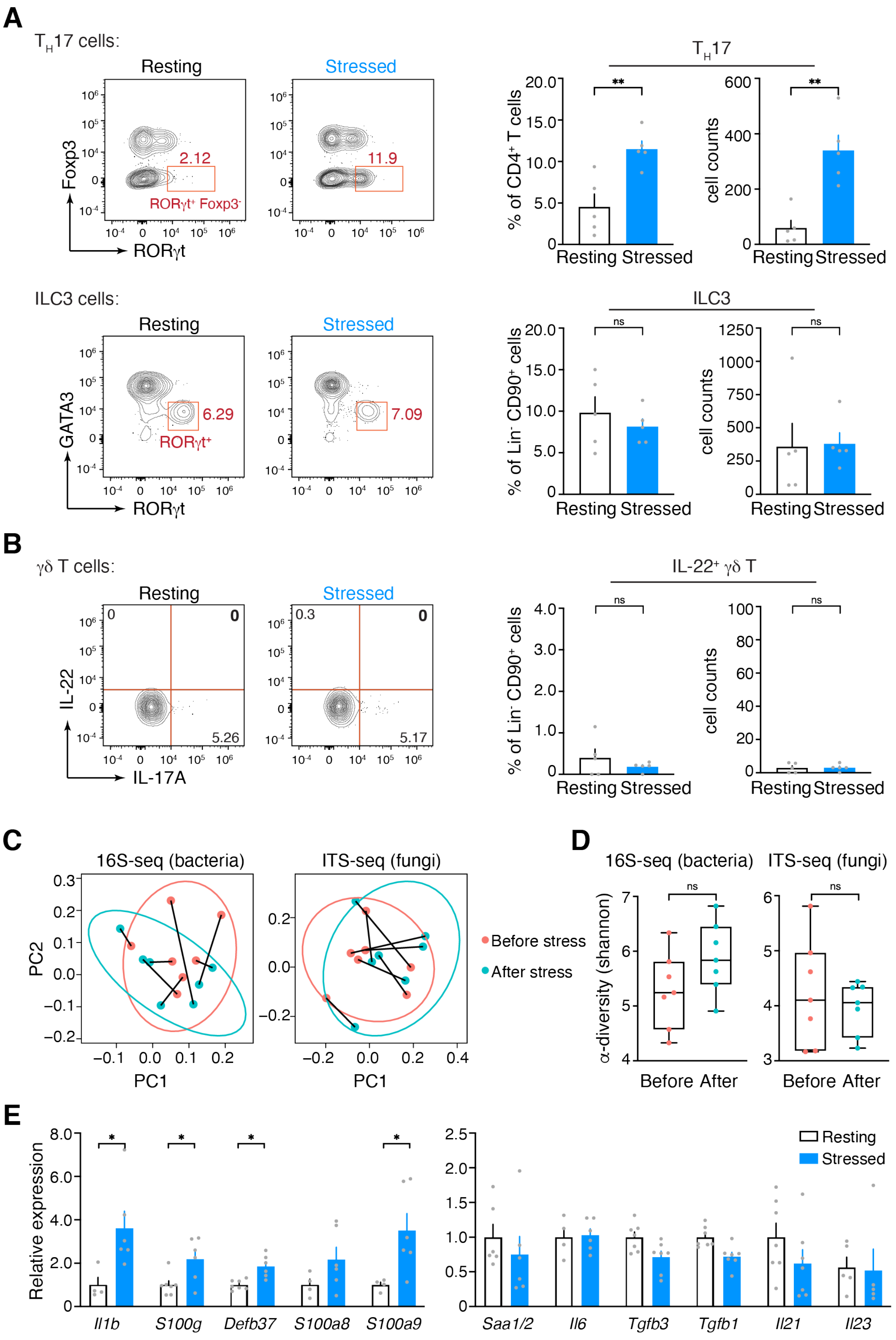

A
TH17 cells:
Resting
Stressed
Foxp3
RORγt
2.12
11.9
RORγt+ Foxp3-
TH17
% of CD4+ T cells
cell counts
Resting Stressed
**
ILC3 cells:
GATA3
6.29
7.09
RORγt+
ILC3
% of Lin- CD90+ cells
ns
B
γδ T cells:
IL-22
IL-17A
0
0.3
5.26
5.17
IL-22+ γδ T
C
16S-seq (bacteria)
ITS-seq (fungi)
PC1
PC2
Before stress
After stress
D
α-diversity (shannon)
Before After
E
Relative expression
Il1b
S100g
Defb37
S100a8
S100a9
Saa1/2
Il6
Tgfb3
Tgfb1
Il21
Il23
*
Resting
Stressed

**Figure S4. Stress leads to immune activation in the intestine.**

(**A,B**) IL-22-producing cells were examined in resting and stressed mice. (A) Intestinal $T_H17$ and ILC3 were examined by RORγt staining and flow cytometry, with representative flow plots shown in the left panel and quantification in the right panel. n= 5 for both groups. (B) Intestinal γδ T cells in resting and stressed mice were examined for IL-22 expression. n= 5 for both groups. (**C,D**) Fecal microbiota analysis of resting and stressed mice. (C) PCoA analysis. 16S-and ITS-sequencing were performed to profile bacteria and fungi, respectively. Samples from individual mice were connected by lines. n=7. (D) Microbiota alpha-diversity was quantified by the Shannon Index. n=7.

(**E**) Expression of intestinal immune genes and $T_H17$-skewing innate immune factors in resting and stressed mice was determined by RT-PCR. n= 4-7 for both groups.

Data are presented as Means ± SEMs. Statistical analysis was performed with the Unpaired t-test (A,B,E) and Paired t-test (D). *P < 0.05; ns, not significant.

Figure S5

A

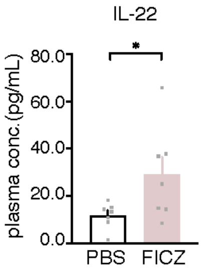


**Figure S5. FICZ treatment increases IL-22 in the blood.**

(**A**) IL-22 concentrations were measured by ELISA after FICZ treatment. n= 7 for both groups. Data are presented as Means ± SEMs. Statistical analysis was performed with the Unpaired t-test (A). *P < 0.05.

**Table S1. Primers used for quantitative RT-PCR in this study.**

| Primer | Sequences |
|---|---|
| *mS100a9*-qF | TGGTGGAAGCACAGTTGGCAAC |
| *mS100a9*-qR | CAGCATCATACACTCCTCAAAGC |
| *mS100a8*-qF | CAAGGAAATCACCATGCCCTCTA |
| *mS100a8*-qR | ACCATCGCAAGGAACTCCTCGA |
| *mS100g*-qF | CTCTCCAAGGAGGAGCTAAAGC |
| *mS100g*-qR | CTCCATCGCCATTCTTATCCAGC |
| *mIl6*-qF | TACCACTTCACAAGTCGGAGGC |
| *mIl6*-qR | CTGCAAGTGCATCATCGTTGTTC |
| *mIl1b*-qF | TGGACCTTCCAGGATGAGGACA |
| *mIl1b*-qR | GTTCATCTCGGAGCCTGTAGTG |
| *mDefb37*-qF | TGCTGCTCCTCTCTCTATCCAA |
| *mDefb37*-qR | TCAGTTTTTGTGGCAACACTTG |
| *mSaa1/2*-rtF | AGTGGCAAAGACCCCAATTA |
| *mSaa1/2*-rtR | GGCAGTCCAGGAGGTCTGTA |
| *mIl21*-rtF | GCCTCCTGATTAGACTTCGTCAC |
| *mIl21*-rtR | CAGGCAAAAGCTGCATGCTCAC |
| *mTgfb3*-rtF | AAGCAGCGCTACATAGGTGGCA |
| *mTgfb3*-rtR | GGCTGAAAGGTGTGACATGGAC |
| *mTgfb1*-rtF | TGATACGCCTGAGTGGCTGTCT |
| *mTgfb1*-rtR | CACAAGAGCAGTGAGCGCTGAA |
| *mIl23*-rtF | CTTCTCCGTTCCAAGATCCTTC |
| *mIl23*-rtF | ACGCACTAGGTTTGCCGAGTAGA |
| *mGapdh*-rtF | AGGTCGGTGTGAACGGATTTG |
| *mGapdh*-rtR | TGTAGACCATGTAGTTGAGGTCA |
